# Metabolic resource overlap impacts on the competition of phyllosphere bacteria

**DOI:** 10.1101/2022.01.20.477054

**Authors:** Rudolf O. Schlechter, Evan J. Kear, Michał Bernach, Daniela M. Remus, Mitja N. P. Remus-Emsermann

## Abstract

The phyllosphere is densely colonised by rich microbial communities, despite sparse and heterogeneously distributed resources. The limitation of resources is expected to drive bacterial competition resulting in exclusion or coexistence based on fitness differences and resource overlap between individual colonisers. We studied the impact of resource competition by determining the effects of different bacterial colonisers on the growth of the model epiphyte *Pantoea eucalypti* 299R (Pe299R). Resource overlap was predicted based on genome-scale metabolic modelling. By combining results of metabolic modelling and pairwise competitions in the *Arabidopsis thaliana* phyllosphere and *in vitro*, we found that ten resources sufficed to explain fitness of Pe299R. An effect of both resource overlap and phylogenetic relationships was found on competition outcomes *in vitro* as well as in the phyllosphere. However, effects of resource competition were much weaker in the phyllosphere when compared to *in vitro* experiments. When investigating growth dynamics and reproductive success at the single-cell resolution, resource overlap and phylogenetic relationships are only weakly correlated with epiphytic Pe299R reproductive success, indicating that the leaf’s spatial heterogeneity mitigates resource competition. Although the correlation is weak, the presence of competitors led to the development of Pe299R subpopulations that experienced different life histories and cell divisions. Surprisingly, in some *in planta* competitions, Pe299R benefitted from the presence of epiphytes despite high resource overlap to the competitor strain suggesting other factors having stronger effects than resource competition. This study provides fundamental insights into how bacterial communities are shaped in heterogeneous environments and provides a framework to predict competition outcomes.

## INTRODUCTION

For bacteria, the leaf surface, i.e., the phyllosphere, is a challenging environment where resources are limited and heterogeneously distributed [1, 2]. However, leaves support bacterial populations of up to 10^7^ CFU per gram of leaf [3]. The number of CFU is impacted by the ability of bacteria to successfully colonise the heterogeneously distributed microenvironments on leaves. These microenvironments thereby influence local interactions and spatial structuring of bacterial communities [4–7].

Microbial communities exhibit intra- and inter-kingdom co-occurrence networks that are shaped and stabilised by priority effects and/or keystone microbial species [8, 9]. The interactions in these networks range from beneficial, neutral, to detrimental, resulting in increased, neutral, or decreased population densities, for at least one of the involved parties, compared to their monoculture [10]. Cooperative or beneficial microbial interactions include cross-feeding, biofilm formation, and cell communication; while competitive or detrimental interactions can be resource competition, contact-dependent antagonism, and secretion of toxic compounds [11]. Competition results in either exclusion or coexistence depending on the differences in the species’ niches and their fitness in an environment [12]. Competitive exclusion is driven by mechanisms that decrease niche differences and increase fitness differences between competing species. Resource use and preference are an important niche axis for a species, and the overlap or similarity with other species is a factor that balances coexistence and competitive exclusion through establishing a level of niche differentiation with others [13].

In the phyllosphere, a large number of bacterial interactions in a community context were shown to be negative [14]. The most parsimonious explanation for the negative interactions is the competition for resources and space, as secretion of antimicrobial compounds appears to be restricted to only a limited number of bacterial taxa in the phyllosphere [15]. Replacement series experiments revealed that pairs of near-isogenic epiphytic bacterial strains with high resource overlap exhibited a strong negative impact on each other’s population size on bean leaves [16]. By contrast, species pairs with a lower resource overlap resulted in larger population sizes than expected [16, 17]. However, this approach is not without its flaws, as it failed to predict competition outcomes between leaf-associated bacteria and a bacterial phytopathogen [18]. A challenge in defining a resource overlap is the lack of information of resource availability, use, and preference of a species in a specific environment. Genome-scale metabolic modelling allows for the study of the metabolic capabilities and nutrient requirements of members within microbial communities in defined growth conditions. Metabolic and community modelling has previously been used in an ecological context to understand the role of metabolic exchange in communities [19], the identifications of keystone species [20], and to define resource overlaps and cross-feeding potentials based on growth requirements [21–23].

As the phyllosphere is highly heterogeneous, ‘coarse-grained’ investigations, such as those considering whole leaves or plants as the units of investigation, are not suited to study local interactions of leaf-associated bacteria. Therefore, to better understand bacterial growth dynamics in the phyllosphere, the micrometre or the single-cell scale must be the resolution of investigation, as every cell may experience a different fate such as microenvironments with different qualities and quantities of nutrients, or competitors [2, 3]. The intimate proximity of bacterial cells should thereby directly impact on community dynamics and short-distance interactions [3, 24, 25].

The CUSPER bioreporter (“repsuc” read backwards, from “reproductive success”), was developed in the epiphytic strain *Pantoea eucalypti* 299R (abb. Pe299R, syn. *Pantoea agglomerans* 299R, *Erwinia herbicola* 299R). Pe299R was originally isolated from a healthy Bartlett pear leaf and has been used in numerous studies since then to understand bacterial physiology and ecology in the phyllosphere [2, 5, 26–29]. Pe299R is part of the order Enterobacterales and the family *Erwiniaceae*. It is a copiotroph, utilises a wide range of nutrients, and grows optimally between 28 and 37°C [17]. CUSPER reports on the number of divisions of individual cells from an initial population based on the dilution of a green fluorescent protein upon cell division and without *de novo* synthesis, and it has shown that Pe299R experience high variations of reproductive success in the phyllosphere [30, 31]. Due to the heterogeneously distributed and limited resources on leaves, the reproductive success of a bacterial cell depends on the local habitability. Consequently, it could be demonstrated that leaves that were pre-colonised with a near-isogenic Pe299R strain reduced the reproductive success of CUSPER bioreporter cells proportional to the pre-coloniser density [32]. However, interspecific competitions at the single-cell resolution in the phyllosphere have not been explored in such detail.

Here, we used genome-scale metabolic modelling to explain competition outcomes in defined growth conditions and in the phyllosphere for phylogenetically diverse leaf-associated bacterial isolates. Resource overlap was determined by metabolic modelling and expected to increase competition, leading to negative impacts on bacterial growth *in vitro* and *in planta*. To that end, the epiphyte *Pantoea eucalypti* 299R (Pe299R) was used as focal strain and challenged with six different phyllosphere-associated bacteria in pairwise competition experiments in different environmental contexts and scales.

## MATERIALS AND METHODS

### Bacterial strains and growth conditions

*Pantoea eucalypti* 299R (Pe299R) and representative epiphytic bacterial strains used in this study are listed in Table 1. Pe299R was used to construct the constitutively red fluorescent protein (mScarlet-I)-producing strain Pe299R::mSc, and was also the parental strain of the CUSPER bioreporter strain Pe299R_CUSPER_ (Pe299R::mSc (pProbe_CUSPER)), which harbours an additional IPTG-inducible green-fluorescent protein gene (Supplemental Materials and Methods). Bacteria were routinely grown on Reasoner’s 2a agar or broth (R2A, HiMedia, India) at 30°C. Minimal media (MM) was used to evaluate growth and competition for defined carbon sources. Minimal media was composed of 1.62 g L^−1^ NH_4_Cl, 0.2 g L^−1^ MgSO4, 1.59 g L^−1^ K_2_HPO4, 1.8 g L^−1^ NaH_2_PO_4_·2H_2_O, with the following trace elements: 15 mg L^−1^ Na_2_EDTA·H_2_O, 4.5 mg L^−1^ ZnSO_4_·7H_2_O, 3 mg L^−1^ CoCl_2_·6H_2_O, 0.6 mg L^−1^ MnCl_2_, 1 mg L^−1^ H_3_BO_3_, 3.0 mg L^−1^ CaCl_2_, 0.4 mg L^−1^ Na_2_MoO_4_·2H_2_O, 3 mg L^−1^ FeSO_4_·7H_2_O, and 0.3 mg L^−1^ CuSO_4_·5H_2_O [33]. Carbon sources used were glucose, fructose, sorbitol, malate, and methanol for determining carbon utilisation profiles and *in vitro* competition assays.

**Table 1.** Phyllosphere-associated bacterial strains.

| Phylum | Phylogroup | PD | Species | Abbr. |
| --- | --- | --- | --- | --- |
| Pseudomonadota | Gammaproteobacteria | n. a. | *Pantoea eucalypti* 299R | Pe299R |
|  |  | 0.41 | *Pseudomonas koreensis* P19E3 | PkP19E3 |
|  |  |  | *Pseudomonas syringae* pv. *syringae* B728a | PssB728a |
|  | Alphaproteobacteria | 0.68 | *Methylobacterium* sp. Leaf85 | MethL85 |
|  |  |  | *Sphingomonas melonis* FR1 | SmFR1 |
| Actinomycetota | Actinobacteria | 0.76 | *Arthrobacter* sp. Leaf145 | ArthL145 |
|  |  |  | *Rhodococcus* sp. Leaf225 | RhodL225 |

Phylogenetic distances (PD), Carbon profile dissimilarities and MRO are in relation to Pe299R. N.a.: Not applicable

### Phylogeny

A phylogenetic tree for the seven phyllosphere-associated strains was constructed based on a multiple sequence alignment of a set of concatenated 31 single-copy genes [34]. Alignment was done with MAFFT, and a phylogenetic tree was then inferred using UPGMA with a Jukes-Cantor model. Concatenation, alignment, and tree inference were performed in Geneious Prime 2022.2.2 (https://www.geneious.com). Newick files were exported into R to retrieve a phylogenetic distance matrix based on branch lengths between strains and Pe299R, using the package *ape* [35].

### *In vitro* growth assays

Each strain was grown at 30°C in R2A broth until the late stationary phase. Cells were then harvested by centrifugation at 2,000 × *g* for 5 min, washed twice in phosphate buffer saline (PBS, 0.2 g L^−1^ NaCl, 1.44 g L^−1^ Na_2_HPO_4_ and 0.24 g L^−1^ KH_2_PO_4_), and resuspended in MM to an optical density at 600 nm (OD_600_) of 0.5. Afterwards, 20 µL of bacterial suspension were added to 180 µL of MM supplemented with a carbon source in flat bottom 96-well microtiter plates (Costar^®^, Corning^®^, NY, USA) with four technical replicates per condition. Carbon utilisation profiles for each species was determined by supplementing MM with a final concentration of 0.2% w/v of a sole carbon source (glucose, fructose, sorbitol, or malate), or 0.2% v/v of methanol. Minimal medium without added carbon source was used as a negative control. The microtiter plates were sealed with a breathable membrane (4ti-0516/96; gas permeability of 0.6 m^3^ m^−2^ day^−1^ and water loss of 1 g m^−2^ day^−1^; Brooks Life Sciences, UK), and incubated at 30°C with shaking. Optical density was measured in a FLUOstar Omega microplate reader (BMG Labtech Ltd., UK) for up to 5 days in 24 h intervals on the same batch culture. For each measurement, ten measurements in different positions of each well were recorded and averaged. The experiments were conducted twice independently. Growth curves of each strain in each growth condition were used to determine growth rate (μ), carrying capacity (K), and area under the curve (AUC), using the *R* package *growthcurver* [36]. These values were used to create a Euclidean distance matrix between species, in which the distance between Pe299R and a second species was used as a proxy of carbon utilisation dissimilarity.

### Construction of genome-scale metabolic and community models

Genomes were retrieved from the PATRIC database, and the annotation files were used to create either individual metabolic models or 2-spp. communities, in which Pe299R was always present, in CarveMe [37]. Models were gap filled using a minimal media composition with (1) carbon sources used for carbon utilisation profiles, or (2) carbon sources detected in *Arabidopsis thaliana* leaves [38, 39]. An index of metabolic resource overlap (MRO) was calculated for each 2-spp. community model using the ‘species metabolic interaction analysis’ (SMETANA) modelling approach developed by Zelezniak *et al*. (2015) [22]. MRO was calculated based on a *in silico* media composition that emulates the growth media tested empirically (Table S1). Similar to the *in vitro* experiments, the composition of the *in silico* media included glucose, fructose, malate, sorbitol, and methanol (M5C). To determine the MRO between pair of species in the phyllosphere, five different media compositions were specified *a priori* based on a range of carbon sources identified in *Arabidopsis thaliana* leaves [38, 39], as the composition of the available resources in the phyllosphere is not yet well defined (L8C, L10C, L13C, L18C, and L26C), detailed in Table S1 [38, 39]. Methanol was included in every media composition, as it is detected on leaf surfaces and is a relevant source of carbon for phyllosphere-associated methylotrophs [40, 41]. Construction of metabolic models and MRO were performed using the High-Performance Computer at ZEDAT, Freie Universität Berlin [42].

### Competition for carbon sources

Competition assays were performed in MM supplemented with mixed carbon sources (MM_5×C_), composed of a total 0.125% w/v of glucose, fructose, sorbitol, malate, and methanol (0.025% w/v glucose, 0.025% w/v fructose, 0.025% w/v malate, 0.025% w/v sorbitol, and 0.025% v/v methanol). The red fluorescent Pe299R::mSc strain was competed against individual non-fluorescent bacteria by mixing both strains in a 1:1 OD_600_ ratio, as described previously [43]. Briefly, flat bottom 96-well microtiter plates (Costar^®^, Corning^®^, NY, USA) were seeded with 200 µL MM_5×C_ containing a defined mixed bacterial suspension (final OD_600_ = 0.05, three technical replicates). The microplate was sealed with a breathable membrane (4ti-0516/96; gas permeability of 0.6 m^3^ m^−2^ day^−1^ and water loss of 1 g m^−2^ day^−1^; Brooks Life Sciences, UK), incubated at 30°C with shaking in a microplate reader, and the red fluorescence of Pe299R::mSc was measured every 5 min for 20 h using an excitation filter at 584 nm and an emission filter at 620-10 nm. Growth parameters from fluorescence growth curves (μ_RFU_, K_RFU_, AUC_RFU_) were determined in *growthcurver*. A competitive ability score based on Chesson’s framework of Coexistence Theory [44] was calculated as in Eq. 1.

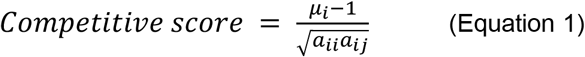

Where *μ*_*i*_ is the growth rate of Pe299R::mSc in monoculture, *a*_*ii*_ is the competition coefficient of Pe299R::mSc when a near-isogenic Pe299R strain is present (intraspecific competition), and *a*_*ij*_ is the competition coefficient of Pe299R::mSc when a different strain is present (interspecific competition). Competition coefficients were calculated as the reciprocal of the corresponding K [45]. For simplicity, competitive ability scores were rescaled and centred to zero (z-score).

### Plant growth

Arabidopsis seeds (*Arabidopsis thaliana* Col-0) were surface-sterilised in a solution containing 50% v/v ethanol and 6% v/v H_2_O_2_ for 90 s, then thoroughly washed three times with sterile distilled water. Before sowing, seeds were stratified in sterile water at 4°C in the dark for at least 2 days. Four seeds were sown aseptically in tissue-culture vessels (Magenta™ GA-7, Magenta LLC., IL, USA) containing 50 mL of ½ Murashige & Skoog (MS; Duchefa, The Netherlands) agar media (1.0% w/v, pH 5.9) in sterile conditions. For gas exchange, the lids of the Magenta GA-7 tissue-culture boxes featured four 1 cm diameter holes that were covered with two layers of 3M Micropore™ tape [46]. Plants were grown in a Conviron A1000 plant growth chamber at 22°C, 80% relative humidity and short day photoperiod (11 h day cycles, light intensity ∼120-150 µE m^-2^ s^-1^).

### Plant inoculation

For plant inoculation, an exponentially growing Pe299R_CUSPER_ culture in lysogeny broth (LB; HiMedia, India) supplemented with 50 µg mL^-1^ kanamycin was induced with 1 mM isopropyl beta-D-1-thiogalactopyranoside (IPTG), as described in detail in Supplemental Material and Methods. Competitor strains were grown on R2A agar plates for 2–5 days, depending on the strain, and a loop of bacteria was resuspended and washed twice in PBS. The IPTG-induced Pe299R_CUSPER_ and the bacterial suspensions were mixed in a 1:1 ratio and adjusted to a final OD_600nm_ of 0.005. Four-week-old arabidopsis plants were inoculated with 200 µL of the bacterial mix per box using a sterile airbrush (KKmoon Airbrush Model T130A). Plants were harvested at 0, 24, 36, and 48 hours post-inoculation (hpi) by cutting the complete leaf rosette from the roots using sterile scissors and scalpel, and transferring the plant into a 1.7-mL microcentrifuge tube. Four individual plants were used per condition. After the fresh weight of each plant was determined, 1 mL PBS with 0.02% v/v Silwet^®^ L-77 was added. Samples were shaken in a bead mill homogenizer (Omni Bead Ruptor 24, Omni International, GA, USA) for two cycles of 5 min at a speed of 2.6 m s^-1^, and sonicated for 5 min (Easy 30 H, Elmasonic, Elma Schmidbauer GmbH, Germany).

Leaf washes were plated onto R2A (total bacterial density) and R2A supplemented with 15 µg mL^-1^ gentamicin (Pe299R_CUSPER_ population), and CFU were determined by serial dilutions and normalised by the corresponding plant fresh weight (CFU gFW^-1^). Growth curve parameters from CFU of Pe299R (μ, K) were used to calculate the competitive ability of an epiphytic strain against Pe299R, as previously described (Eq. 1). The remaining supernatants were transferred into a sterile 1.7-mL microcentrifuge tube and centrifuged at 15,000 × *g* for 10 min at 4°C to collect cells for microscopy. Cells were resuspended and fixed in 100 µL of fixative solution (4% w/v paraformaldehyde –PFA– in PBS) overnight at 4°C. After this period, cells were washed twice in PBS and resuspended in 20 µL PBS. Then, one volume of 96% v/v ethanol was added to the samples. Samples were stored at -20°C until further analysis.

### Microscopy

Cells recovered from leaves after 0, 24, and 36 hpi were mounted on microscopy slides coated with 0.1% w/v gelatine. Images were acquired on a AxioImager.M1 fluorescent widefield microscope (Zeiss) at 1000× magnification (EC Plan-Neofluar 100×/1.30 Ph3 Oil M27 objective) equipped with the Zeiss filter sets 38HE (BP 470/40-FT 495-BP 525/50) and 43HE (BP 550/25-FT 570-BP 605/70), an Axiocam 506 (Zeiss), and the software Zen 2.3 (Zeiss). At least 100 cells were acquired per biological replicate in three different channels: green (38HE filter set), red (43HE filter set), and phase contrast.

### Image analysis

Images were analysed in FIJI/ImageJ v. 2.0.0-rc-69/1.52s [47]. As Pe299R_CUSPER_ constitutively expresses mScarlet-I, the red fluorescent channel was used as a mask to select individual cells, using the thresholding method “intermodes” and converted into a binary mask object. Only particles in a size range of 0.5–2.5 µm were considered, excluding cells on the edges of the field of view. All objects were manually inspected using the phase contrast images to corroborate the selection of bacterial cells, and to exclude false positive red fluorescent particles. The mask was then used to determine green fluorescence of Pe299R_CUSPER_ cells in the green-fluorescent channel to calculate the reproductive success (RS) of single cells. In addition, background fluorescence was measured by sampling a random section of background area in each fluorescent image [2].

### Estimation of single-cell reproductive success

The RS of the Pe299R_CUSPER_ bioreporter is calculated as the number of divisions a GFP-loaded cell underwent after arrival to a new environment. This estimation is based on the dilution of the green-fluorescent signal after cell division, as *de novo* biosynthesis of GFP is transcriptionally repressed [30]. The RS of Pe299R_CUSPER_ cell at a time *t* was estimated from background-corrected fluorescence measurements by subtracting the mean background fluorescence from the mean fluorescence intensity of each cell in each field of view. Then, the reproductive success of a cell *n* at time *t* (RS_n,t_)—number of divisions of an immigrant cell since its arrival to a new environment—was calculated as

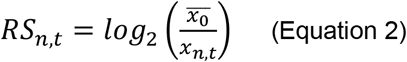

Where 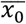 is the mean intensity of the cell population at time zero, and *x*_*n,t*_ the fluorescence intensity of a single cell *n* at time *t* [31].

As the Pe299R_CUSPER_ bioreporter decreases in intensity upon each cell division, the RS value from the background intensity measurements in the green-fluorescent channel was calculated to define a limit of detection (LOD) for Pe299R_CUSPER_. The LOD was defined as the RS value that has a 5% probability of being background noise. Consequently, calculated values of RS for single cells above this threshold were grouped, as the number of generations that a cell with low fluorescent intensity underwent cannot be further estimated. The distribution of RS from the initial cell population was determined as a relative fraction of Pe299R_CUSPER_ cells from the total observed population, by binning cells into subpopulations with different RS values: RS_0_: RS < 0.5; RS_1_: 0.5 ≤ RS < 1.5; RS_2_: 1.5 ≤ RS < 2.5; RS_3_: 2.5 ≤ RS < 3.5; RS_4_: 3.5 ≤ RS < 4.5; RS_>4_: RS ≥ 4.5. Non-Metric Multidimensional Scaling (NMDS) and Permutational Multivariate Analysis of Variance (PERMANOVA) with Bray-Curtis dissimilarities was selected to evaluate the variation of a Pe299R_CUSPER_ population (as relative fractions) explained by multivariate data (i.e., time of sampling and presence of an epiphyte). Bray-Curtis dissimilarity matrix and PERMANOVA with 999 permutations were performed using the *vegan* package [48].

### Data analysis

If not stated otherwise, all data processing, statistical analyses, and graphical representation were performed in R [49]. Data processing and visualisation were performed using the *tidyverse* package [50]. Graphical representations of matrices were constructed using the *ComplexHeatmap* package [51]. Pearson’s correlations (*r*) were used to compare variables using the *cor()* function of the *stats* package. Linear or generalised linear regressions were constructed with the *lm()* or *glm()* function from the *stats* package, respectively, to evaluate the effect of the presence of an epiphyte, metabolic resource overlap (MRO), and/or time of sampling with the competitive ability of a strain against Pe299R in different environments (*in vitro* and in the phyllosphere). ANOVA were performed using the *aov()* function of the *stats* package. Eta squared (η^2^) was used to measure the effect size of the predictors in the regression models using the *lsr* package [52].

## RESULTS

### Differences in carbon utilisation are predicted by genome-scale metabolic modelling

Differences in growth on different carbon sources and the metabolic resource overlap (MRO) between the focal species *Pantoea eucalypti* 299R (Pe299R) and members of actinobacteria, gamma-, and alphaproteobacteria (Fig. 1) were determined empirically and based on genome-scale metabolic models, respectively.

**Figure 1.**
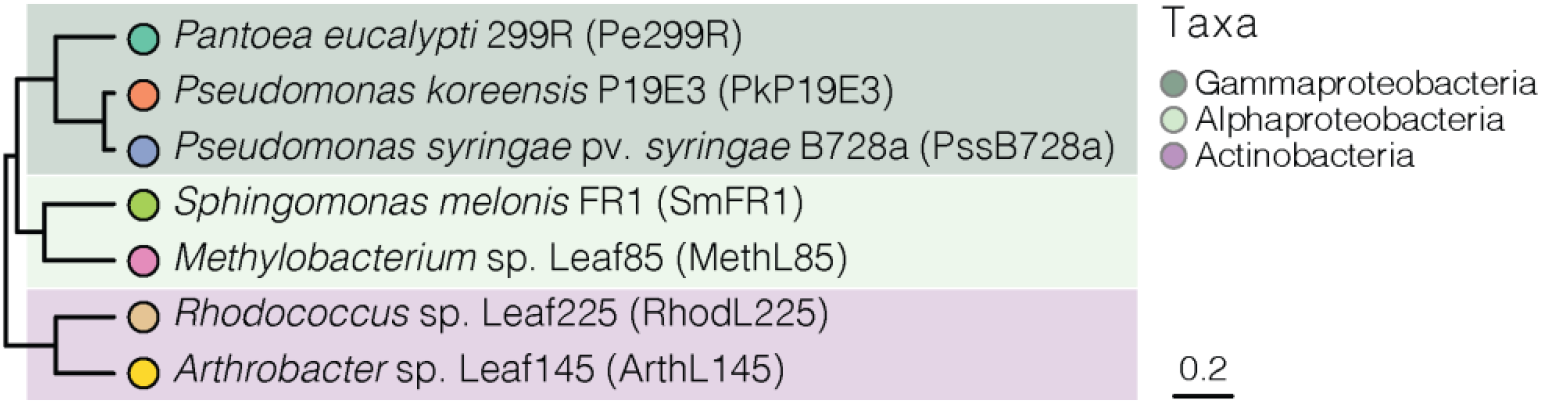
Phylogenetic tree of phyllosphere-associated bacterial strains. An UPGMA tree was created for the strains used in this study using a set of 31 single-copy marker genes [34]. Scale bar represents the number of substitutions per site.

Individual strains were grown in MM supplemented with either 0.2% w/v glucose, fructose, malate, sorbitol, or 0.2% v/v methanol (Fig. S1a). Growth rate (μ) and carrying capacity (K) were retrieved for each growth curve and used to cluster the strains based on similarity (Fig. 2a, Table S2). Hierarchical clustering based on utilised resources placed Pe299R in a clade with the gammaproteobacterium *Pseudomonas koreensis* P19E3 (PkP19E3) and the actinobacterium *Arthrobacter* sp. Leaf145 (ArthL145). These strains were able to grow on glucose, fructose, and malate, reaching high and similar K in liquid media. *Sphingomonas melonis* FR1 (SmFR1) and *Pseudomonas syringae* pv. *syringae* B728a (PssB728a) showed a similar utilisation pattern as the first clade. However, SmFR1 population did not reach a similar K, while PssB728a was also able to grow in sorbitol. The most dissimilar strains in relation to Pe299R were *Rhodococcus* sp. Leaf225 (RhodL225) and *Methylobacterium* sp. Leaf85 (MethL85). The resulting carbon utilisation profile was used as an empirical distance matrix for resource use dissimilarity between Pe299R and a second strain.

**Figure 2.**
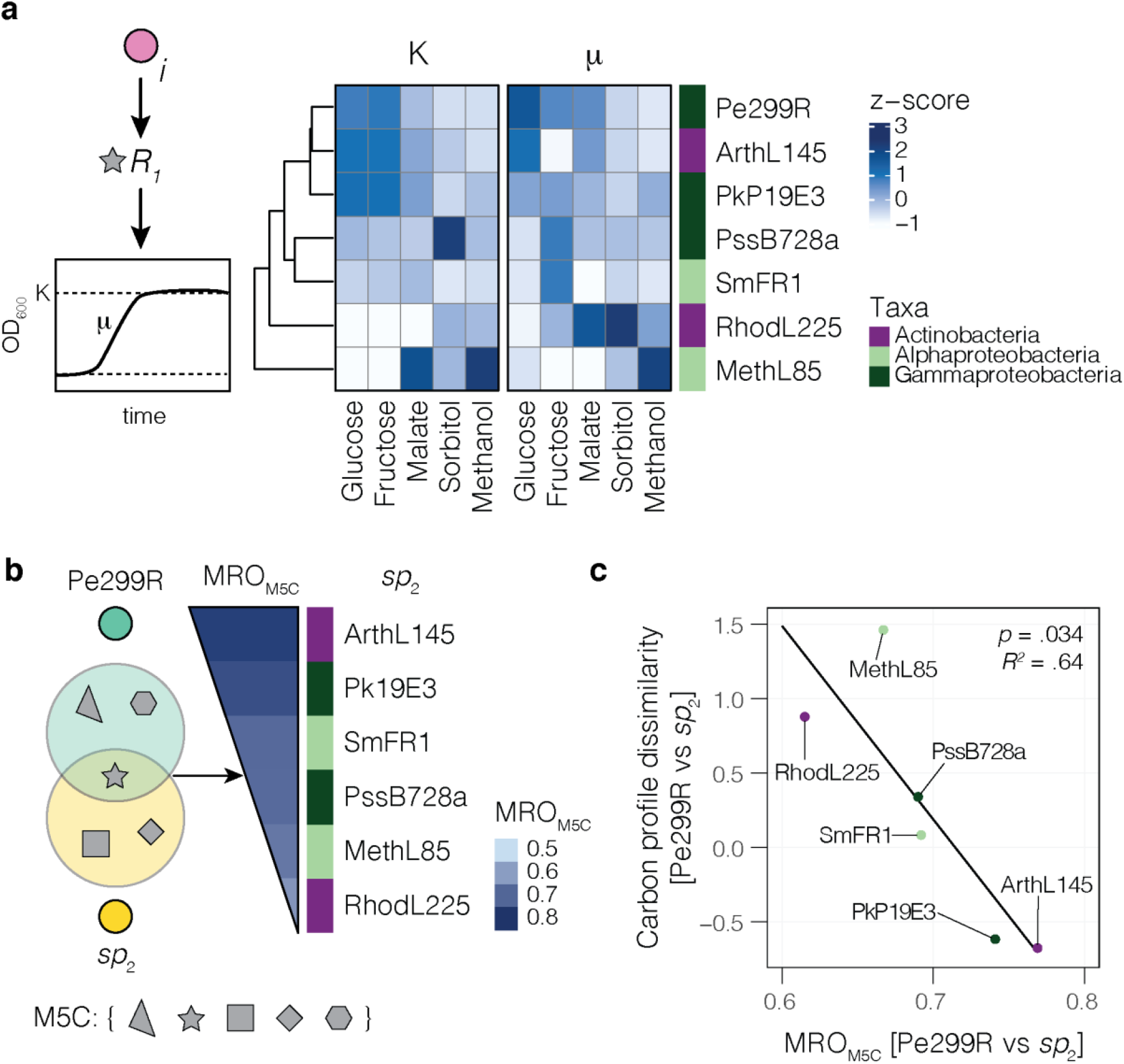
Metabolic resource overlap portrays differences in empirical carbon utilisation profiles between Pe299R and a second strain. **(a)** Carbon utilisation matrix. Bacterial strains were clustered based on carrying capacity (K) and growth rates (μ) from growth in minimal medium supplemented with individual carbon sources. Values were rescaled into z-scores for hierarchical clustering. **(b)** Metabolic resource overlap (MRO) is an index of resource similarity modelled, based on genomic information, under *in silico* media composition including glucose, fructose, malate, sorbitol, and methanol (M5C). Strains were ranked based on descending MRO_M5C_ with Pe299R. **(c)** Linear relationship of MRO_M5C_ and carbon profile dissimilarity between Pe299R and a second strain (*sp*_2_).

The MRO is an estimation of the maximal overlap between the minimal growth requirements of two (or more) metabolic models [22]. Thus, MRO calculates the potential of species to compete for a list of compounds defined *a priori*. From a list that includes glucose, fructose, malate, sorbitol, and methanol as carbon sources (M5C), MRO_M5C_ was calculated for Pe299R and secondary strains (Fig. 2b, Table 2). The strain pair Pe299R–ArthL145, as well as Pe299R–PkP19E3 showed the highest MRO_M5C_ values (0.77 and 0.74, respectively), which were part of the same cluster based on empirical growth profile, while the lowest MRO_M5C_ and highest profile dissimilarity were observed between the pairs Pe299R– MethL85 and Pe299R–RhodL225 (MRO_M5C_ of 0.67 and 0.62, respectively). The carbon profile dissimilarity between Pe299R and a second strain was strongly correlated with MRO_M5C_ (*r* = -0.85, *p* = 0.032) but not with phylogenetic distances (*r* = 0.27, *p* = 0.60). Additionally, MRO_M5C_ was a predictor of carbon profile dissimilarity between the focal and other strains (Fig. 2c). Linear regression analysis showed a negative relationship between MRO and carbon profile dissimilarity (R^2^ = 0.65, *F*_1,4_ = 10.41, *p* = 0.032), suggesting that the use of genome-scale metabolic modelling can be used to explain strain differences in resource use in a defined environment.

**Table 2.** Dissimilarity metrics and competition scores of phyllosphere-associated strains in relation to Pe299R.

| Strain | *in vitro* |  |  | *in planta* |  |  |  |  |  |
| --- | --- | --- | --- | --- | --- | --- | --- | --- | --- |
|  | Carbon profile dissimilarity | MRO<sub>M5C</sub> | Competition score | MRO<sub>L8C</sub> | MRO<sub>L10C</sub> | MRO<sub>L13C</sub> | MRO<sub>L18C</sub> | MRO<sub>L26C</sub> | Competition score |
| PkP19E3 | 1.89 | 0.74 | 2.64 | 0.64 | 0.64 | 0.64 | 0.64 | 0.69 | -0.75 |
| PssB728a | 4.02 | 0.69 | 3.03 | 0.59 | 0.67 | 0.59 | 0.59 | 0.77 | 1.31 |
| MethL85 | 6.52 | 0.67 | 0.92 | 0.69 | 0.69 | 0.62 | 0.69 | 0.69 | 0.81 |
| SmFR1 | 3.45 | 0.69 | 1.05 | 0.64 | 0.64 | 0.62 | 0.62 | 0.88 | -1.03 |
| ArthL145 | 1.75 | 0.77 | 2.41 | 0.69 | 0.81 | 0.85 | 0.81 | 0.81 | 0.51 |
| RhodL225 | 5.22 | 0.62 | 0.69 | 0.69 | 0.62 | 0.69 | 0.69 | 0.77 | -0.86 |

Carbon profile dissimilarities and MRO are in relation to Pe299R

### Competition *in vitro* is driven by resource overlaps

To confirm the predictions of the genome-scale metabolic modelling, the ability of a competitor to affect the growth of a fluorescently red-labelled Pe299R strain (Pe299R::mSc) was evaluated *in vitro*. First, the optical density of every strain was measured in MM supplemented with multiple resources (MM_5×C_: 0.025% w/v glucose, 0.025% w/v fructose, 0.025% w/v malate, 0.025% w/v sorbitol, and 0.025% v/v methanol) to confirm that each strain was able to growth under these conditions (Fig. S1b). To test the effect of a strain on the growth of Pe299R, Pe299R::mSc was mixed in a 1:1 ratio with a second strain and red fluorescence intensity was measured over time (Fig. 3a). If normalised by the fluorescence signal of a monoculture, constitutive fluorescence expression was shown to serve as a proxy for changes in bacterial biomass of individual strains in pairwise competitions [43].

**Figure 3.**
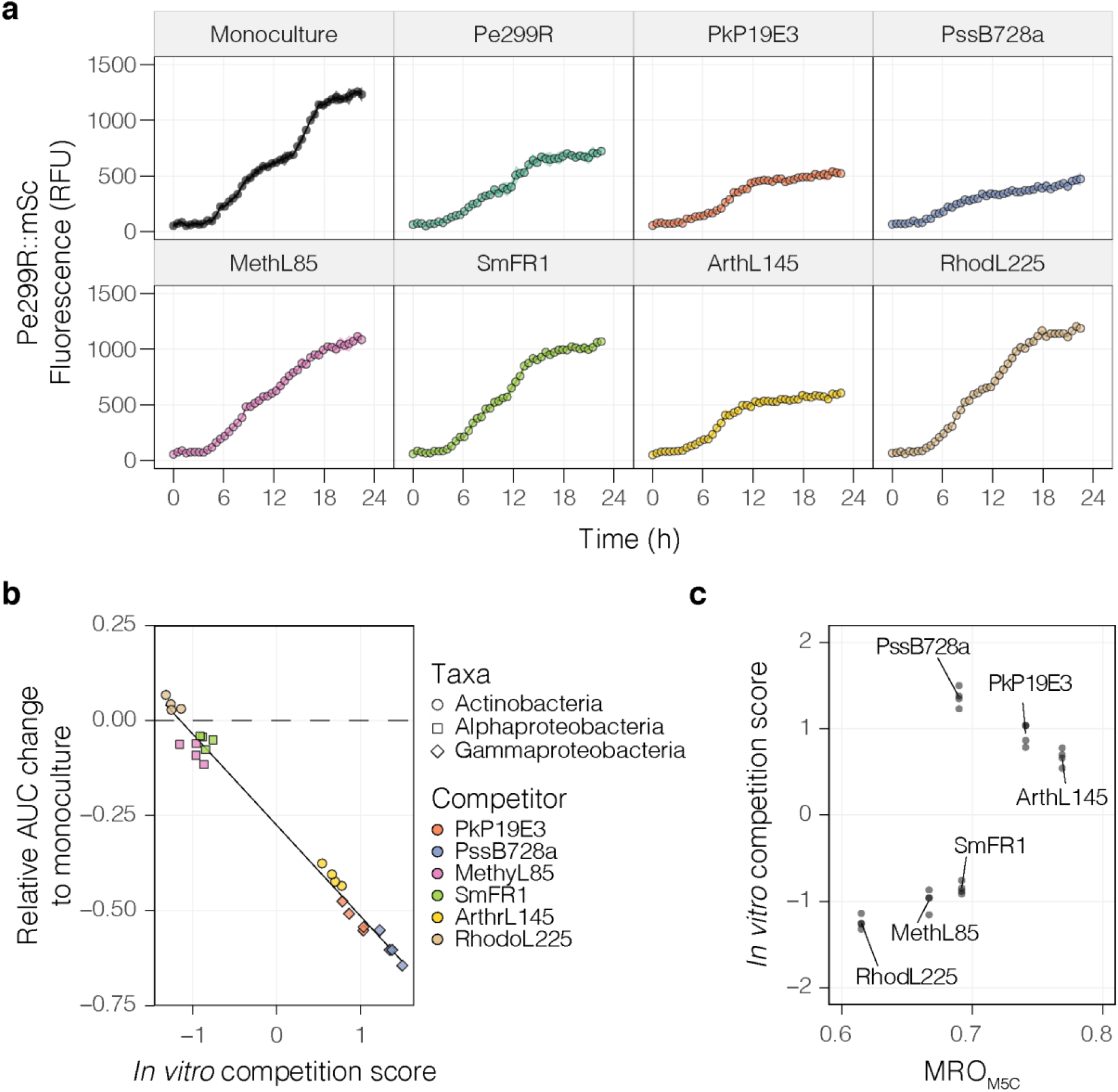
Pe299R is affected by the presence of a competitor *in vitro*. **(a)** Fluorescence curves of Pe299R::mSc co-inoculated with a competitor (top label) in MM_5×C_. **(b)** Relationship between competition score (Eq. 1) and the relative change in the area under the fluorescent curve of Pe299R::mSc in the presence of a competitor in relation to the monoculture. **(c)** Relationship between MRO_M5C_ and the competitive score of a second epiphyte against Pe299R. Details of the regression model can be found in Table S4.

Growth parameters from the fluorescence curves (μ_RFU_, K_RFU_, AUC_RFU_) were retrieved and compared with a competitive ability score (Eq. 1, Table S3). This competition score includes both μ_RFU_ and K_RFU_ in interspecific competition (Pe299R::mSc vs. *sp*_2_) in comparison to intraspecific competition (Pe299R::mSc vs Pe299R) and the monoculture (Pe299R::mSc). The competition score showed a strong correlation with most metrics (|*r*| > 0.96), except with growth rate alone (Fig. S2, *r* = 0.67). Thus, changes in Pe299R growth in relation to the monoculture can be explained by the ability of a strain to compete with Pe299R (Fig. 3b, R^2^ = 0.98, *F*_1,25_ = 1272, *p* < 0.05). The highest competition scores were observed for PssB728a, PkP19E3, and ArthL145, while the lowest were RhodL225, MethL85, and SmFR1. Regression analysis was used to evaluate the effect of MRO and/or phylogenetic distances in the competition scores against Pe299R (Table S4). These competition outcomes were partially explained by MRO_M5C_ (Fig. 3c, R^2^ = 0.46, *F*_1,22_ = 20.67, *p* < 0.05). PssB728a showed the largest deviation from the regression model, suggesting that mechanisms other than competition for carbon could explain the increased competitive ability of PssB728a *in vitro*. However, no interference competition was observed in double-layer assays on R2A (Fig. S3). By excluding PssB728a from this analysis, MRO_M5C_ became a strong predictor of competitive ability (R^2^ = 0.81, *F*_1,18_ = 83.45, *p* < 0.05). Alternatively, a generalised linear model including MRO_M5C_, phylogenetic distance (PD), and the interaction between these terms explained the competitive ability of an epiphyte against Pe299R (Table S4, Gamma error distribution with a log link, pseudo-R^2^ = 0.89, *F*_1,20_ = 9.61, *p* = 0.0056). In this model, the competitive score of an epiphyte depends on the interaction between MRO_M5C_ and PD (MRO_M5C_ × PD: *p* = 0.0062), in which the competitiveness of closely-related species to Pe299R are less dependent on MRO_M5C_ than distantly-related species (Fig. S4a). These results indicate that the *in vitro* competitiveness of an epiphyte against Pe299R in a defined medium can be explained by the utilised resources that they have in common, as predicted by the MRO, and their phylogenetic distance.

### Bacterial competition in the phyllosphere at different scales reflect different competition outcomes

#### Competition at the population scale

The effect of a competitor on the growth of Pe299R was evaluated in the arabidopsis phyllosphere by co-inoculating four-week-old arabidopsis plants with Pe299R_CUSPER_ (Pe299R::mSc (pProbe_CUSPER)) and a second strain to estimate changes in population densities as well as single-cell reproductive success of Pe299R *in planta*.

In every case, total bacterial density was determined and increased over time to a similar maximal load (Fig. S5). Particularly, changes in CFU of Pe299R were dependent on both sampling time and presence of a competitor (Time × Competitor: *F*_1,21_ = 3.34, *p* < 0.05). Compared to monoculture, only the presence of SmFR1 and the near-isogenic Pe299R strain negatively impacted the Pe299R_CUSPER_ population at 24 and 48 hpi, respectively (Fig. 4a).

**Figure 4.**
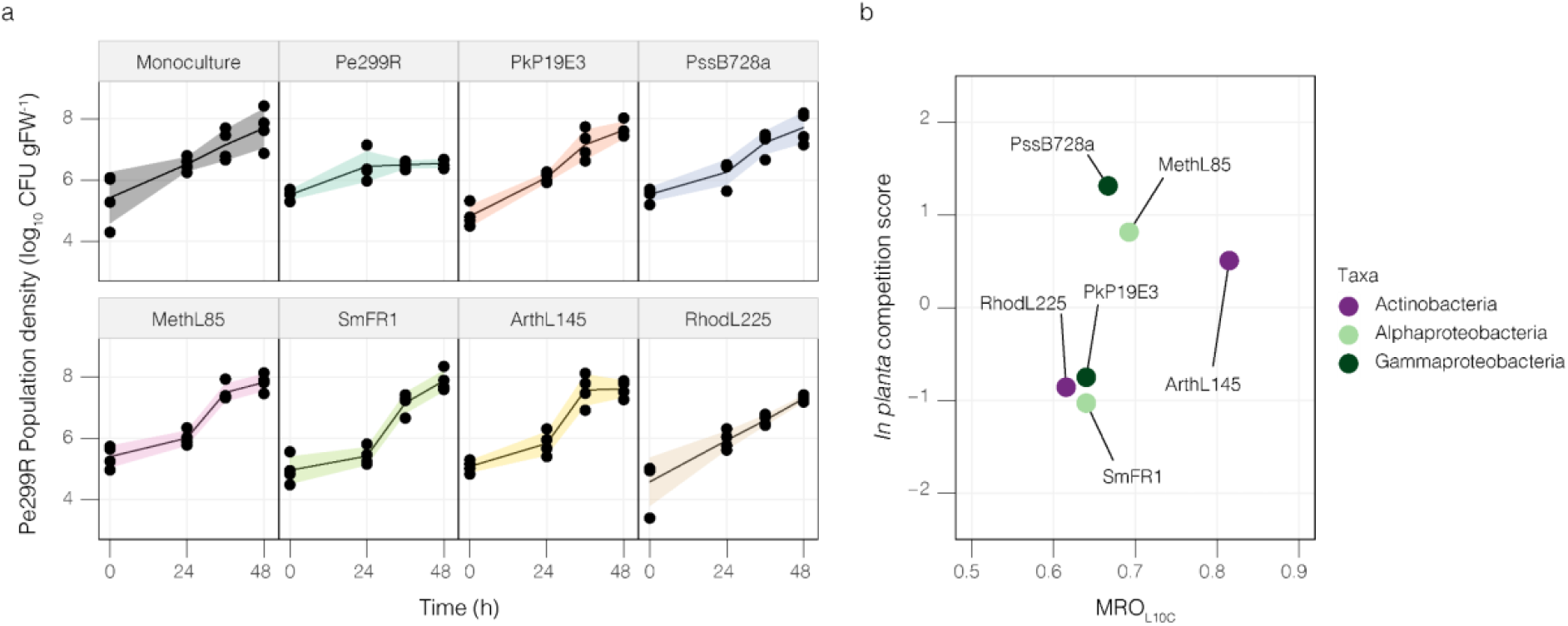
Changes of Pe299R population density in the phyllosphere. **(a)** Population size of *P. eucalypti* 299R::Tn*7*::mSc::Gm^R^(pProbe_CUSPER) (*Pe*299R_CUSPER_) on arabidopsis plants as monoculture or in the presence of a second epiphyte (top label). Each data point represents the CFU of Pe299R per gram of fresh leaf weight (CFU gFW^-1^) of individual plants (n = 4) at different sampling points (0, 24, 36, and 48 h). Groups were compared using two-way ANOVA, with a significance level of α = 0.05. **(b)** Relationship between resource overlap (MRO_L10C_) and the competition score of an epiphyte against Pe299R in the phyllosphere. Details of the regression model can be found in Table S5.

Although the presence of a competitor did not largely affect the Pe299R_CUSPER_ population at the CFU-level, different MRO indexes were calculated based on carbon sources that have been detected on arabidopsis leaves in an effort to explain differences in competitive abilities in this environment (Fig. S6, Table 2). The MRO with the most predictive power was the one calculated from a medium composition including ten carbon sources (MRO_L10C_): fumarate, sucrose, aspartate, malate, citrate, glutamate, alanine, fructose, threonine, and methanol (L10C, Table S1). These resources were the ten most abundant metabolites detected in arabidopsis leaves [39]. Particularly, compared to a similar composition including eight resources (L8C, Table S1), the presence of citrate, alanine, and threonine, as well as the absence of glucose, increased the predictability of competition outcomes, through an increase in the MRO between Pe299R and ArthL145 (MRO_L8C_ = 0.69; MRO_L10C_ = 0.81), and Pe299R and PssB728a (MRO_L8C_ = 0.59; MRO_L10C_ = 0.67). However, this effect was significant only when a linear regression model included MRO_L10C_ and the phylogenetic distance (PD) between an epiphyte and Pe299R (Fig. 4b, R^2^ = 0.92, *F* = 20.15, *p* = 0.048). The regression model suggests that the competitive ability of an epiphyte against Pe299R depends on both their resource overlap and phylogenetic relationships (Table S5, MRO_L10C_ × PD: *p* = 0.029). In the phyllosphere, high competition scores were observed among closely related species with high MRO_L10C_ (Fig. 4b, Fig. S4b). In summary, Pe299R population density in the phyllosphere was not largely affected by the presence of a competitor, and differences in the competitive ability of this second strain could be explained by both its resource overlap and phylogenetic relationship with Pe299R.

#### Competition at the single cell-resolution

An improved version of the CUSPER bioreporter plasmid was constructed and was used to develop Pe299R_CUSPER_ (Fig. S7). In contrast to the initial CUSPER bioreporter, Pe299R_CUSPER_ constitutively expresses a red fluorescent protein and carries the recently developed green fluorescent protein mClover3 in a multicopy plasmid, rather than a chromosomally inserted single copy of GFPmut3. Pe299R_CUSPER_ is a bioreporter that estimates the reproductive success (RS) of immigrant cells in a new environment by back-calculating the number of divisions a cell underwent since its arrival [30–32].

The RS was determined by measuring the reduction in single cell green fluorescence compared to the mean green fluorescence of the population at time zero (*t*_0_), *ex situ*. Thereby, the reproductive success of a population and individual cells can be estimated. The limit of detection was determined based on the empirical cumulative distribution function from background fluorescence signals (Fig. S8a). A 5% probability represents RS values equal or greater than 4.58. Thus, a limit of detection of 4.5 cell divisions was selected (Fig. S8b). Cells with RS values above 4.5 were grouped and considered to undergo more than four divisions (RS_>4_).

The relative increase in Pe299R_CUSPER_ population from the initial inoculum at a given time of sampling can be estimated based on the fraction of cells in a particular subpopulation and the number of divisions that a cell is expected to undergo upon arrival in the phyllosphere [31]. The increase in Pe299R_CUSPER_ population from single-cell data was associated with the increase in population size at the CFU level (R^2^ = 0.63, *F*_1,382_ = 655.6, *p* < 0.05), suggesting that the single-cell measurements and the threshold used were adequate to assess changes in Pe299R_CUSPER_ populations. Similar to the results of the CFU-based population-level experiment above, the presence of competitors did not affect the average single-cell reproductive success of the Pe299R_CUSPER_ population compared to the monoculture (Fig. S9).

The distribution of immigrant cells that experienced different levels of reproductive success was analysed as relative fractions of the initial population. Cell groups were binned based on the number of divisions that the respective ancestral immigrant cell underwent after inoculation. The different bins for the number of divisions –the reproductive success– ranged from 0 to >4 generations after inoculation. Consequently, a population structure of Pe299R_CUSPER_ was defined based on the relative fraction of cells with different RS. The variation in the population structure of Pe299R_CUSPER_ could be explained by the time of sampling (R^2^ = 0.10, *F*_1,55_ = 8.69, *p* = 0.001) and the presence of competitors (R^2^ = 0.27, *F*_7,55_ = 3.36, *p* = 0.002) using PERMANOVA (Table S6). The initial population was composed of cells with zero (RS_0_) or one division (RS_1_) across the different treatments, and were excluded from the multivariate analysis, as it only accounts for the initial population and not for underlying competitive interactions. At 24 and 36 hpi, cells that divided three and more times contributed most to the final populations (Fig. 5). For Pe299R_CUSPER_ in the presence of Pk19E3, PssB728a, ArthL145, RhodL225, and MethyL85, a relatively high fraction of cells that divide between zero and three times was observed at 24 hpi, whose distribution became bimodal (Fig. S10). However, only the presence of PssB728a (R^2^ = 0.24, *F*_1,14_ = 4.50, *p* = 0.011) and MethL85 (R^2^ = 0.36, *F*_1,14_ = 7.80, *p* = 0.003) led to a differentiation in the population structure of Pe299R_CUSPER_ compared to the monoculture (Fig. S11). Compared to Pe299R_CUSPER_ as monoculture, the presence of PssB728a and MethL85 increased the relative fraction of Pe299R_CUSPER_ cells that divided more than four times. As PssB728a and MethL85 exhibited an MRO_L10C_ with Pe299R of 0.67 and 0.69, respectively, which lie within one standard deviation from the mean MRO_L10C_ of the tested strains (0.68 ± 0.072), the effect of epiphytes on the structure of the Pe299R population in the phyllosphere cannot be associated solely with their resource overlaps.

**Figure 5.**
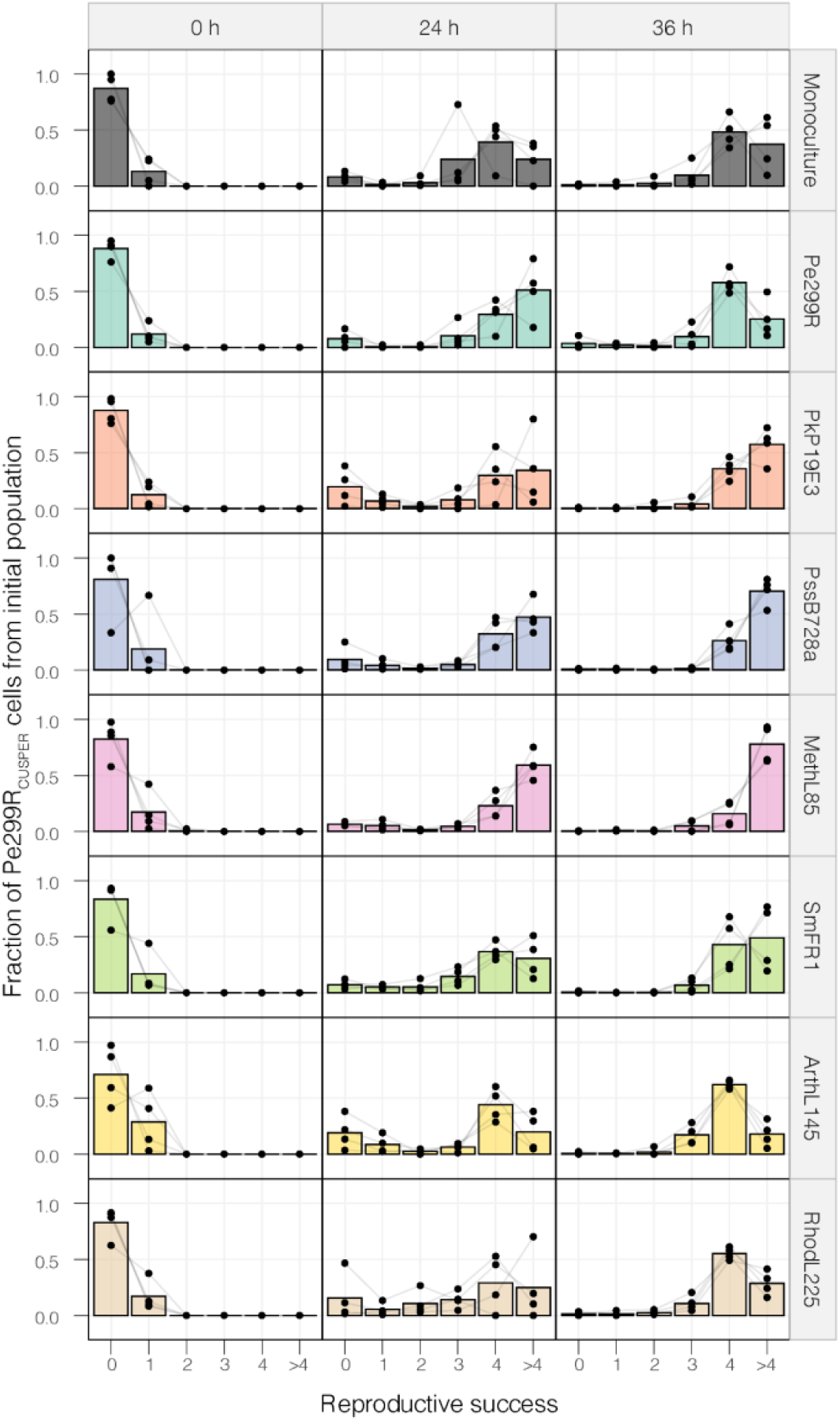
Changes in the composition of the Pe299R_CUSPER_ populations over time in the presence of a second epiphyte. Relative fraction of the reproductive success of the founder population to the observed Pe299R_CUSPER_ population at each sampling time point (0, 24, and 36 hpi) as monoculture or in the presence of a second epiphyte. The relative fractions of each biological replicate are shown and connected with a grey line. The bar represents the mean relative fraction for all replicates.

## DISCUSSION

Understanding microbial community structure and dynamics in the phyllosphere requires a deeper investigation into the mechanisms that influence local microbe-microbe interactions. As resources are a limiting factor for bacterial colonisation [53], and negative interactions are common outcomes between microbes in the phyllosphere [14], we hypothesised that resource competition is the dominant type of interaction in this environment. A negative correlation between coexistence and similarity in resource utilisation has been shown for a number of pairs of epiphytic bacteria, including the focal species Pe299R [17]. On arabidopsis, plant-protective *Sphingomonas* spp. decrease the population of the phytopathogen *Pseudomonas syringae* pv. *tomato* DC3000. Although some sphingomonads suppress the proliferation of *P. syringae* pv. *tomato* DC3000 via priming of the plant immunity [54], the high resource overlaps between *P. syringae* pv. *tomato* DC3000 and the *Sphingomonas* spp. suggest that resource competition explains in parts the decrease in population size of the pathogen [55]. However, estimating bacterial resource preferences in complex and heterogeneous environments is challenging. In this work, genome-scale metabolic modelling was used to predict the outcome of species interactions under homogeneous *in vitro* conditions as well as in the heterogeneous phyllosphere.

Similarity in resource utilisation has been used to define a niche overlap index that includes a wide range of carbon sources, many of which are unlikely to be relevant on the leaf surface [17]. The availability of resources in a given environment and resource preference of competitors determine the effective resource overlap [56]. MRO is an index that incorporates the minimal growth requirements of a species’ metabolic model under defined media composition in silico [22]. To our knowledge, MRO has not been used in combination or validated with empirical studies. Our results show that MRO reflects the dissimilarities in resource utilisation in bacterial batch cultures. To test whether MRO also predicts competition outcomes in heterogeneous environments, it was selected to link the similarity of resource preferences to competitive differences among epiphytes on leaves. Similar to other natural environments, the resource landscape on leaves is uneven and otherwise challenging to measure [1, 2, 57, 58]. The MRO calculated from the ten most abundant resources in the arabidopsis leaf metabolome [39] were predictive for the competition outcomes of the here-studied strains. It is worth noting that the selected resources cannot be generalised for competitions on leaves, as other strains could compete for resources that were not included in the MRO calculation, such as low abundant or rare resources [59], vitamins [60], or iron [57, 61]. However, MRO calculated using additional resources did not increase the explanatory power of resource overlap in this study, suggesting that major competition in the phyllosphere is restricted to a limited set of most abundant resources.

Generally, predictability of the competition ability *in vitro* and, especially in the phyllosphere improved when phylogenetic distances were accounted for as a factor in the analysis. Phylogenetic distance was included in the model, as taxon-dependent characteristics may favour either high or low phylogenetic diversity [62]. For example, competition between *Pseudomonas fluorescens* SBW25 and other species decreased at increasing phylogenetic distances, which correlated with increased niche differences [63]. However, MRO does not exclusively correlate with phylogenetic distances, which is congruent with previous findings showing that genes associated with carbon source utilisation are not phylogenetically conserved [64]. The lack of phylogenetic conservation also holds true for plant-associated bacteria, as comparative genomic analysis of the arabidopsis microbiota showed a high overlap of genes linked to carbon and amino acid metabolism, independent of their phylogeny [65]. The results presented in this work suggest that evolutionary-conserved traits contribute to competition outcomes in the phyllosphere. Traits such as aggregation, motility, communication, production of biosurfactants and/or siderophores could influence the fitness of leaf colonisers. Our results are therefore congruent with modern coexistence theory, where competitive exclusion depends on niche differences (*e*.*g*., high resource overlap) and fitness differences between competing species [12, 66].

Pe299R_CUSPER_ was used to measure the single-cell reproductive success of Pe299R and to estimate bacterial fitness in competition *in planta*. This bioreporter relies on the fluorescence intensity of individual cells, which can be traced back to the dilution of a fluorescent protein after cell division [30]. The number of divisions that can be determined is however limited to the initial four cell divisions. This bioreporter was instrumental in understanding that bacterial populations in the phyllosphere separate into subpopulations over time [30].

The observed RS heterogeneity within arabidopsis-colonising Pe299R_CUSPER_ is congruent with findings in the phyllosphere of bush bean leaves (*Phaseolus vulgaris*) [30]. This supports the notion that variable habitability is a common feature of the phyllosphere of different species. The plant host impacts on bacterial colonisation, suggesting that the host could influence bacteria-bacteria interactions by environment modifications through variations of metabolite availability during the circadian cycle [38], leaf side [67], leaf development [68, 69], ageing [70], and cuticle composition [71]. Differences in reproductive success within the Pe299R population correlate with, but are not limited to, spatially distinct resource pools, such as carbohydrates and water, on leaves [1, 2, 72]. Considering the variable fate of bacterial cells during leaf colonisation, the effect of resource overlap in bacterial interactions was expected to be evaluated at the single-cell level. While we showed that the presence of other strains did not lead to differences compared to the Pe299R_CUSPER_ monoculture, PssB728a and MethL85 positively affected RS at the single cell level.

Despite showing the highest population-level competitive scores while not featuring a notably low MRO, the fraction of successful Pe299R cells (>4 divisions) were higher in the presence of PssB728a compared to the monoculture. This suggests that mechanisms other than resource competition were influencing the interactions between Pe299R and PssB728a. The pseudomonad PkP19E shows a similar, albeit not statistically significant, effect by increasing the fraction of cells in the Pe299R population that has a RS >4. *Pseudomonas* spp. produce biosurfactants, i.e. amphiphilic molecules that decrease water surface tension and thereby increase resource permeability onto the leaf surface, and increase bacterial survival due to their water retaining hygroscopic nature [73–76]. By producing biosurfactants, *Pseudomonas* spp. could thereby benefit Pe299R. Alternatively, these strains could engage in cooperative interactions such as cross-feeding, as observed between *Pantoea* spp. and *Pseudomonas koreensis* in the *Flaveria robusta* leaf apoplast [77]. However, further investigations are required to understand the mechanisms that result in beneficial interactions in the phyllosphere.

MethL85 belongs to a group of resource specialists and facultative methylotrophs from the genus *Methylobacterium*. Methanol utilisation is a fitness advantage in the phyllosphere, as methylotrophs can utilise the released methanol from the plant cell wall metabolism [40]. As one carbon metabolism is highly overrepresented in proteomes of methylobacteria on leaves [78, 79], it is expected that MethL85 utilises methanol as a main carbon source and does not compete with Pe299R for their preferred carbon sources. Possibly, additional biomass and the biosurfactant production of the strain may act as a water retaining factor which increases survival and spread of bacteria [73, 74, 80]. Hence, additional growth of Pe299R in presence of MethL85 compared to a near isogenic co-inoculant is not unexpected.

Our results suggest that there is little impact of resource overlap on the competition between bacteria that co-colonise the leaf surface. This could be a result of the strong segregation of habitable sites on leaves and low initial bacterial densities at the time of inoculation. Thus, resource competition in combination with historical contingency caused by priority effects could have a larger impact on competition outcomes. Although co-colonisation had little effect on Pe299R, this could change after pre-emptive colonisation by a competitor [32, 61].

Overall, we observed a relationship between the resource overlap and the competition of pairs of species in both homogeneous and heterogeneous environments. This relationship was stronger and more predictive *in vitro* compared to the phyllosphere. However, single-cell measurements did not correlate with population-level measurements, indicating that competition is operating at the micrometre, or single-cell resolution and thus, local competition cannot be investigated by measuring interactions and changes in population densities at the whole-leaf scale. Regardless, our findings support an important role of resource overlap in community assembly processes of bacteria in the phyllosphere. This is in line with previous findings that related resource overlap of competitors with disease severity on tomato plants [81] and co-existence of near-isogenic strains that differed only in the ability to metabolise an additional resource [17].

Understanding the impact of resource competition during bacterial community assemblage in the phyllosphere has major implications in developing effective biocontrol strategies against phytopathogens [82, 83]. Many bacterial foliar pathogens undergo an epiphytic phase during the initial colonisation of leaves [84]. This phase is characterised by population growth before invading the endophytic compartments. The rational design of biocontrol agents or communities to reduce pathogen populations in the phyllosphere through competitive interactions could prevent crop losses caused by microbial diseases. However, our findings showed that colonisation prevention of leaves by bacteria that feature different degrees of resource overlap in the phyllosphere is challenging. Previous metrics of resource overlap that considered many different resources are not the best strategy to select for strong competitors against a focal species [17]. Instead, resource overlap metrics should consider resources that are most relevant in the system and for the phytopathogen to be controlled, and traits that are phylogenetically conserved. In this study, ten resources detected in arabidopsis leaves had the most predictive power. Additional resources did not increase the predictive outcome of the metric (Table S5). Thus, we consider MRO in conjunction with information of resource abundances in the phyllosphere of arabidopsis more suitable than previous metrics.

## ACKNOWLEDGEMENTS

We would like to thank Paula Jameson and Matthew Stott for insightful discussions. This work was supported by Marsden Fast Start grant number 17-UOC-057 (M.N.P.R.-E.). R.O.S. was supported by a New Zealand International Doctoral Research Scholarship (NZIDRS) and a University of Canterbury College of Science Ph.D. scholarship. M.B. was supported by a University of Canterbury Ph.D. scholarship. We would also like to thank the HPC Service of ZEDAT, Freie Universität Berlin, for computing time.

