## Supplemental Material for "Metabolic resource overlap impacts on the competition of phyllosphere bacteria"

**Preprint Servers:** biorxiv

**Author Contributions:** R.O.S. and M.N.P.R-E. designed research; R.O.S., E.J.K., M.B., D.M.R. performed research; R.O.S analysed data; and R.O.S., and M.N.P.R-E. wrote the paper.

**Competing Interest Statement:** The authors declare no conflict of interest.

This file includes:

Extended Materials and Methods

Supplemental Figures S1 to S12

Supplemental Tables S1 to S7

References

### Extended Materials and Methods

#### Cloning of pProbe_CUSPER

The plasmid pProbe_CUSPER (Fig. S7) was constructed via isothermal assembly [1]. To that end, DNA fragments were obtained through polymerase-chain reaction (PCR) using Phusion High-Fidelity DNA polymerase (Thermo Scientific), following the manufacturer's recommendations. Annealing temperatures ($T_a$) were chosen based on the respective melting temperature ($t_m$) of the primers (Table S7). Touchdown PCRs were performed to amplify PCR products with overlapping ends for isothermal assemblies, as described elsewhere [2]. The *lac* promoter $P_{A1/04/03}$ was amplified using as template the plasmid miniTn7(Gm)PA1/04/03-eyfp-a [3], which was a gift from Tim Tolker-Nielsen (Addgene plasmid # 111620), and the primers Plac_fw and Plac_rv (Table S7). The green fluorescent protein gene, *mClover3*, was amplified from the plasmid pMRE147 [2] using the primers fp_fw and fp_rv. The repressor element *lacI*$^{q}$ was amplified from the plasmid pCPP39 using the primers lacIq_fw and lacIq_rv. Finally, the gentamicin resistant gene *gmR* was amplified from the plasmid pMRE143 using the primers gmR_fw and gmR_rv. Following amplification, PCR fragments were purified using Monarch PCR clean up kit (New England Biolabs, Place). The plasmid pFru97 [4] was used for cloning of the pProbe_CUSPER backbone by restriction digest using HindIII and ScaI, which includes ori*V*, *rep*, *mob*, a terminator region and the *nptII* gene as selection marker. Fragments were consolidated into pProbe_CUSPER and used to transform Stellar$^{TM}$ Competent Cells (*E. coli* HST08, Takara Bio Inc., Japan).

#### Development of the *Pe*299R$_{\text{CUSPER}}$ bioreporter

The constitutively red fluorescent protein-expressing strain *Pantoea eucalypti* 299R::Tn7::mScarlet-I::Gm$^{R}$ (Pe299R::mSc) [2] was transformed with pProbe_CUSPER (Fig. S7) through electroporation to obtain the new bioreporter *Pe299R*::Tn*7*::mScarlet-I::Gm$^{R}$(pProbe_CUSPER), from here onwards referred to as Pe299R$_{\text{CUSPER}}$. Pe299R$_{\text{CUSPER}}$ is an updated version of the CUSPER bioreporter [5], in that every genetic material is harboured in the plasmid pProbe_CUSPER and that it encodes the fluorescent protein mClover3, a brighter version than GFPmut3. Electrocompetent cells were prepared based on Gonzales *et al.* (2013) [6] with modifications. Overnight culture of Pe299R::mSc grown on nutrient agar (NA, HiMedia, India) supplemented with 15 µg mL$^{-1}$ gentamicin was scraped off with a loop and resuspended in phosphate buffer saline (PBS, 0.2 g L$^{-1}$ NaCl, 1.44 g L$^{-1}$ $Na_2HPO_4$ and 0.24 g L$^{-1}$ $KH_2PO_4$). The bacterial suspension was washed in cold 10% v/v glycerol twice by centrifugation (5 minutes at 5,000 × *g*). Finally, cells were resuspended in 100 µL 10% v/v glycerol and transferred into an ice cold 2-mm electroporation cuvette. For electroporation, 100 ng of plasmid pProbe_CUSPER were added into an ice cold 2-mm cuvette (Genesee Scientific) and mixed carefully. One pulse of 1.8 kV at 200 Ω and 25 µF was applied, then 900 µL of nutrient broth (NB) was quickly added and the mix was transferred to a 15-mL conical tube. Cells were incubated for 2 h at 30 °C with shaking (200 rpm), and then plated onto NA supplemented with 50 µg mL$^{-1}$ kanamycin (Km). Pe299R$_{\text{CUSPER}}$ was routinely grown on NA with Km at 30°C.

#### IPTG optimisation

Expression of mClover3 in pProbe_CUSPER is under the control of the *lac* promoter $P_{A1/04/03}$ and repressed by LacI$^{q}$, which can be inhibited by the addition of isopropyl beta-D-1-thiogalactopyranoside (IPTG). To determine the optimal concentration of IPTG for mClover3 expression, fluorescence was measured at different concentrations of IPTG in a FLUOstar Omega microplate reader (BMG Labtech). Pe299R$_{\text{CUSPER}}$ and the parental Pe299R::mSc strain were grown until the mid-exponential phase ($OD_{600nm}$ ~ 0.5) in NB. Cells were collected by centrifugation (5 min at 2,000 × *g*) and washed twice with PBS. Bacterial suspensions were adjusted to an $OD_{600}$ of 0.5 and 20 µL were used to seed each well of a flat bottom 96-well microtiter plate (Costar) containing NB supplemented with increasing concentrations of IPTG (0-5 mM). Fluorescence intensity was measured in 15 min intervals in a FLUOstar Omega microplate reader (BMG Labtech) for 20 h (Fig. S12). mClover3 fluorescence was measured using an excitation filter of 485-12 nm and an emission filter of 540-10 nm, while OD was detected with a 600 nm filter.

#### Growth inhibition

To determine potential interference competition between focal species and Pe299R, a double layer assay was conducted as described elsewhere [7, 8]. In brief, overnight culture of Pe299R::mSc in R2A was harvested by centrifugation (5,000 × *g* for 5 min), washed twice in PBS, and mixed in soft R2A agar (0.75% w/v agar) kept at 40°C with a temperature-controlled magnetic stirrer (MR Hei-Tec, Heidolph) to reach a concentration of $OD_{600}$ 0.01 (top layer). This solution was quickly poured onto an agar plate containing 15 mL of R2A (bottom layer). Then, fresh cultures of each focal species grown on R2A agar were resuspended, washed twice in PBS, and adjusted to an $OD_{600}$ of 1.0. The top layer was drop-inoculated with 2 µL of each bacterial suspension and plates were incubated at 30 °C for 2 days. As control, a 2 µL drop of 15 mg $mL^{-1}$ gentamicin and 5 mg $mL^{-1}$ tetracycline were deposited onto the top layer to inhibit Pe299R growth locally. The experiments were conducted twice independently with three technical replicates in each experiment.

### SUPPLEMENTAL FIGURES

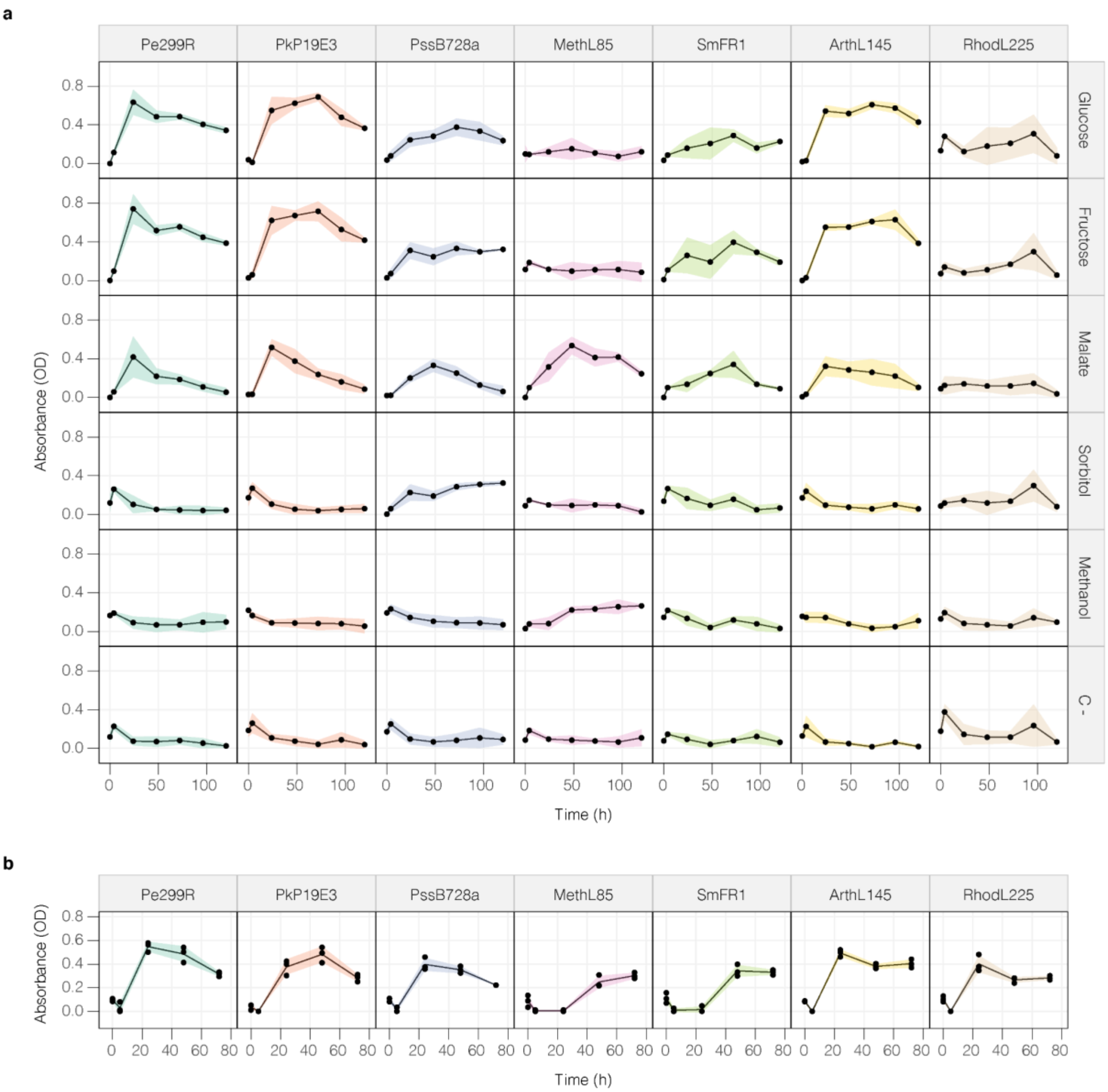


**Figure S1. Growth of epiphytic bacteria in different carbon sources. (a)** Growth of epiphytic bacteria in minimal media supplemented with 0.2% w/v of glucose, fructose, malate, sorbitol, 0.2% v/v methanol. **(b)** Growth of epiphytic bacteria in minimal media supplemented with 0.125% w/v of mixed carbon sources ($MM_{5×C}$). Growth was measured as the absorbance at 600 nm ($OD_{600}$) over time. As negative control, MM without carbon sources was used. Coloured area indicates the area covered by the mean ± SD at each time point.

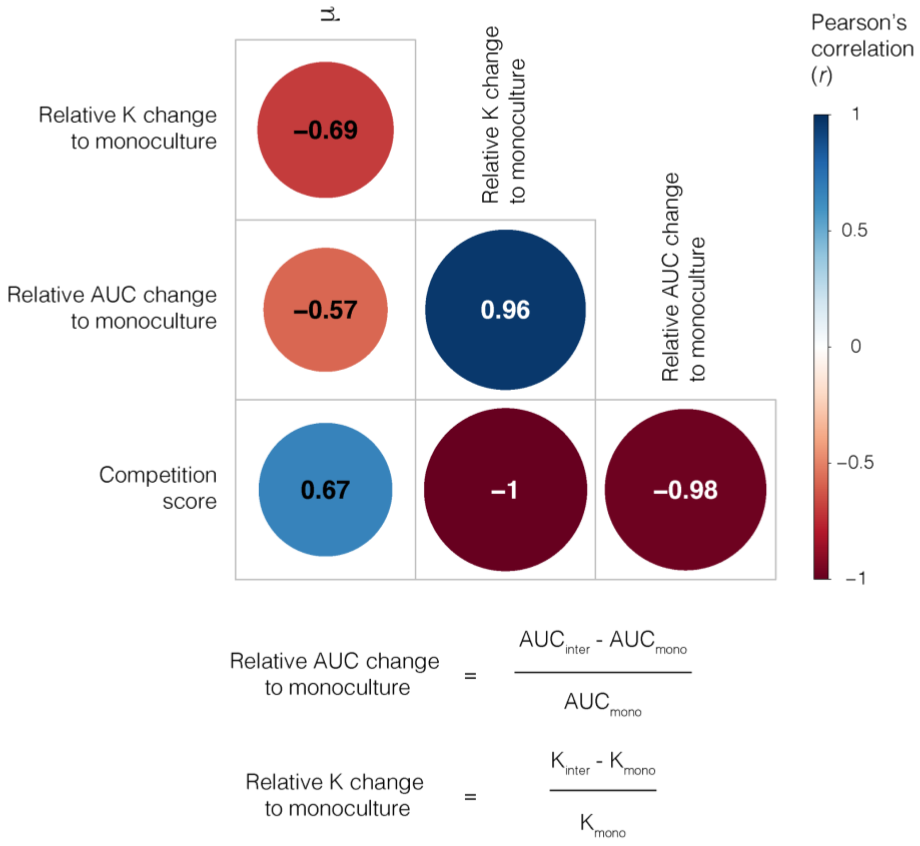


**Figure S2. Comparison of competitive ability score of epiphytes and Pe299R fluorescence curve parameters.** Pearson's correlation coefficients (*r*) between competition score (Eq. 1) of an epiphyte against Pe299R in $MM_{5\times C}$ and the growth rates (μ), relative change in carrying capacity (relative K change), and relative change in area under the fluorescent curve (relative AUC change) of Pe299R in the presence of an epiphyte in relation to the monoculture. Relative change to monoculture was calculated as the fractional difference between K or AUC of Pe299R in interspecific competition in relation to Pe299R as monoculture.

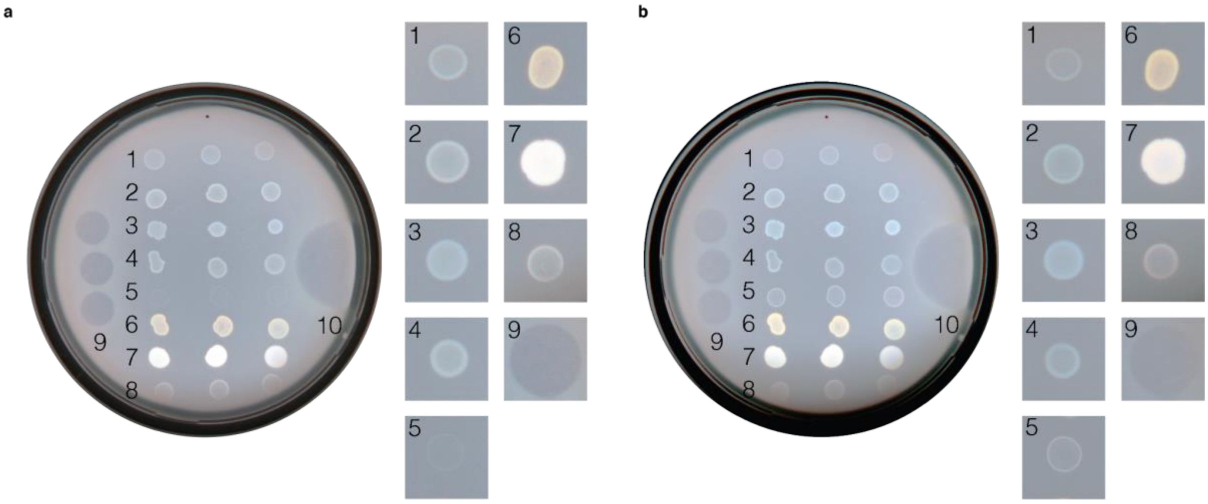


**Figure S3. Growth inhibition assay.** Growth of bacterial strain onto a top layer containing *Pantoea eucalypti* 299R ($OD_{600}$ = 0.01) in R2A at **(a)** 24 and **(b)** 48 h of incubation at 30 °C. In **(a)** and **(b)**, the top layer was inoculated with: (1) Pe299R::mSc; (2) Pe299R; (3) PkP19E3; (4) PssB728a; (5) MethL85; (6) SmFR1; (7) ArthL145; (8) RhodL225; (9) Gentamicin; (10) Tetracycline.

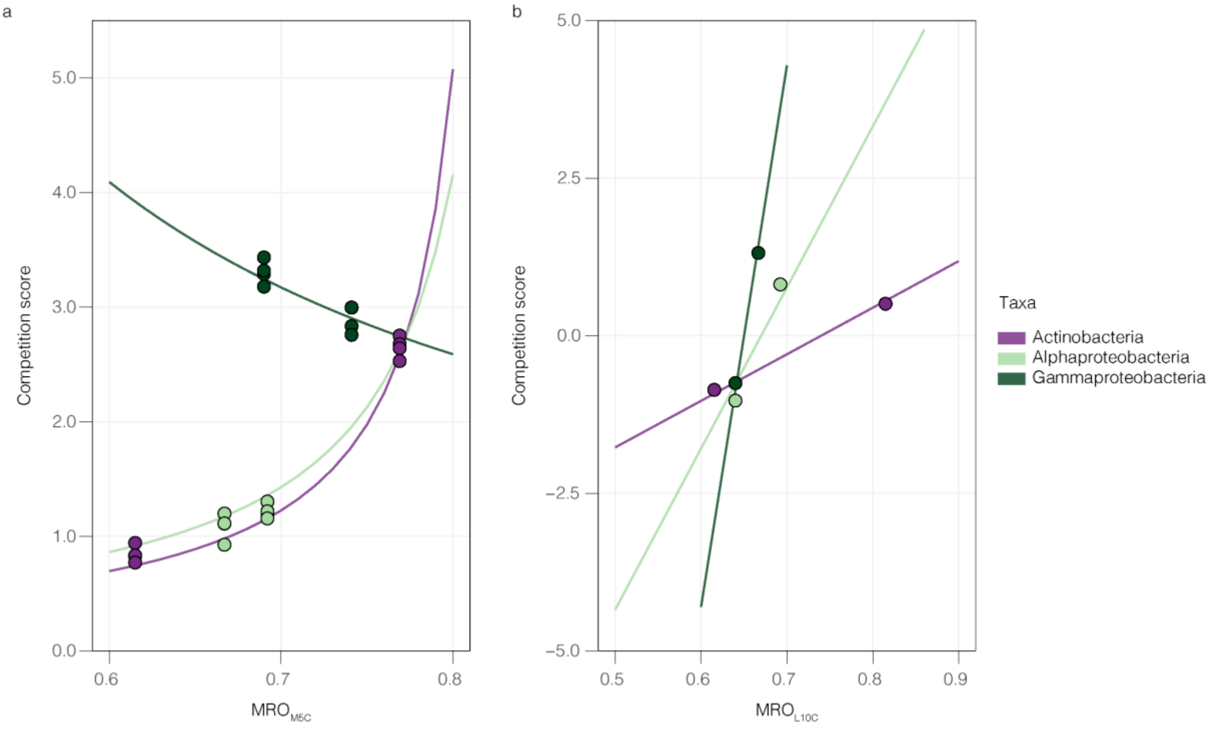


**Figure S4. Resource overlap and phylogenetic distance influence competition scores of epiphytes against Pe299R.** Regression models of competition scores explained by MRO and the phylogenetic group of an epiphyte in relation to Pe299R in **(a)** minimal medium (M5C) and in **(b)** the phyllosphere (L10C). Details of the regression models are found in Table S4 and Table S6.

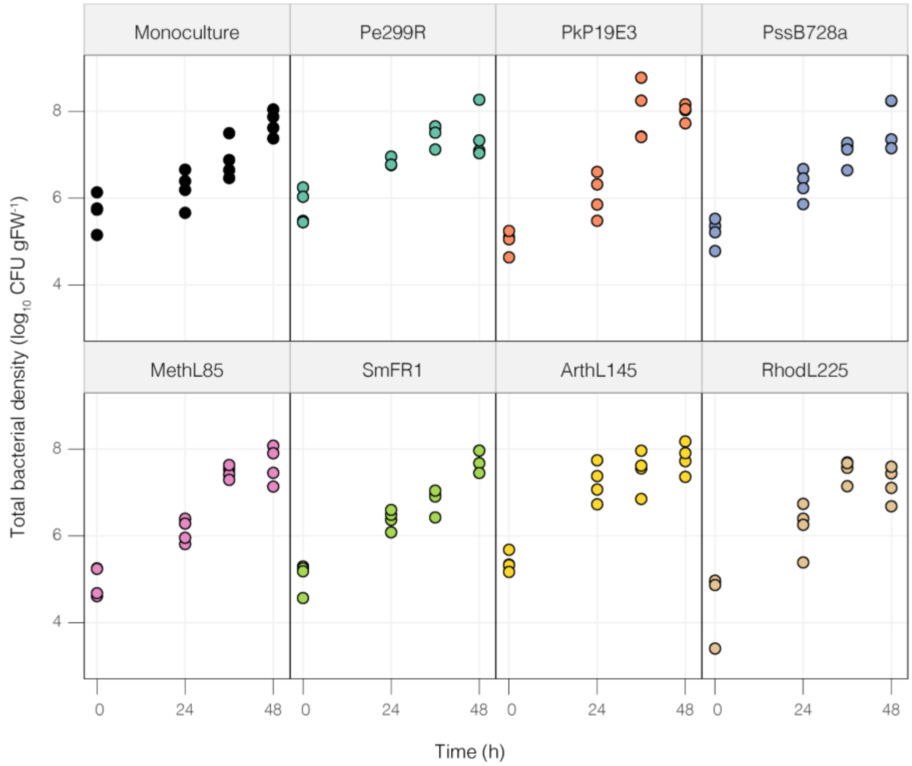


**Figure S5. Total bacterial density in the phyllosphere**. Each data point represents the total CFU counts per gram of fresh leaf weight (CFU $gFW^{-1}$) of independent plants (n = 4) at different sampling points (0, 24, 36, and 48 h). In each plot, Pe299R was co-inoculated with a second epiphyte (top label).

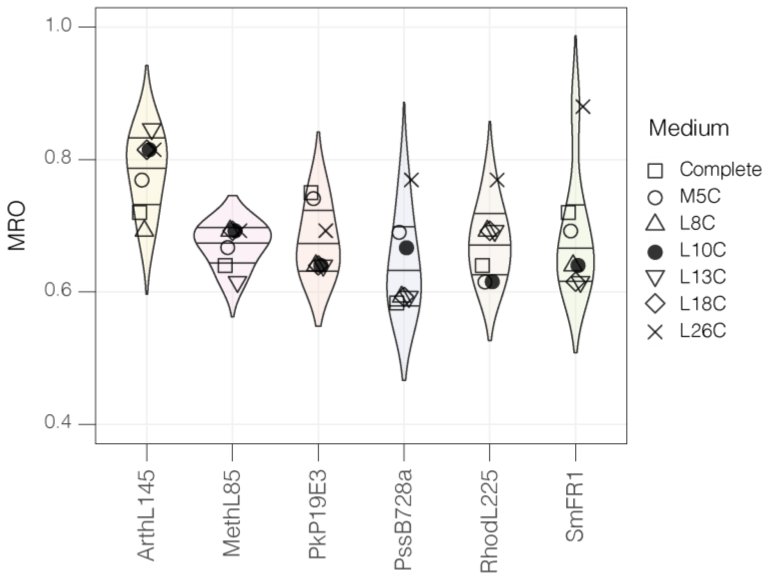


**Figure S6. MRO between Pe299R and a second epiphyte in different growth compositions.** MRO values from 2-spp. communities including Pe299R and a second epiphyte calculated from different environments. L10C indicated as a filled point is the medium whose MRO showed the most predictive power. A detailed list of each *in silico* medium composition can be found in Table S5.

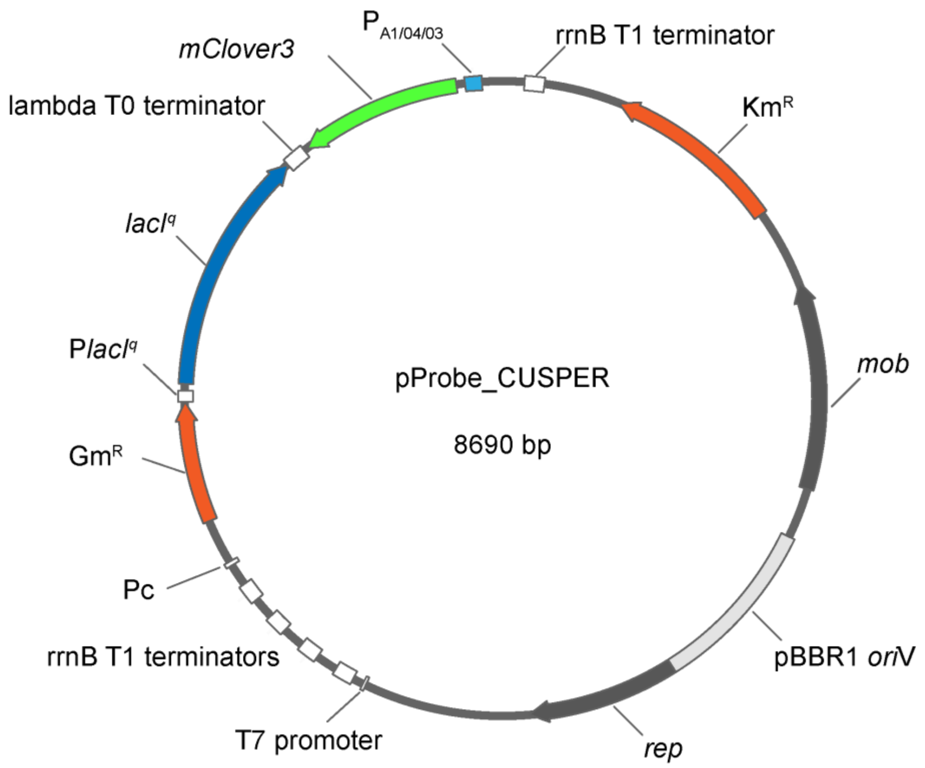


**Figure S7. pProbe_CUSPER plasmid map.** Plasmid-borne CUSPER bioreporter (8,690 bp). Green fluorescent protein gene variant, *mClover3*, is under the control of the *lac* promoter $P_{A1/04/03}$. Selection markers are a kanamycin resistance (Km$^{R}$) and a gentamicin resistance (Gm$^{R}$) gene. Origin of replication pBBR1 *oriV*. Genetic elements include repressor protein gene *lac*$^{Iq}$, replication protein gene, *rep*, and mobilisation protein genes, *Mob*. Other genetic elements such as terminators and promoters are indicated in white.

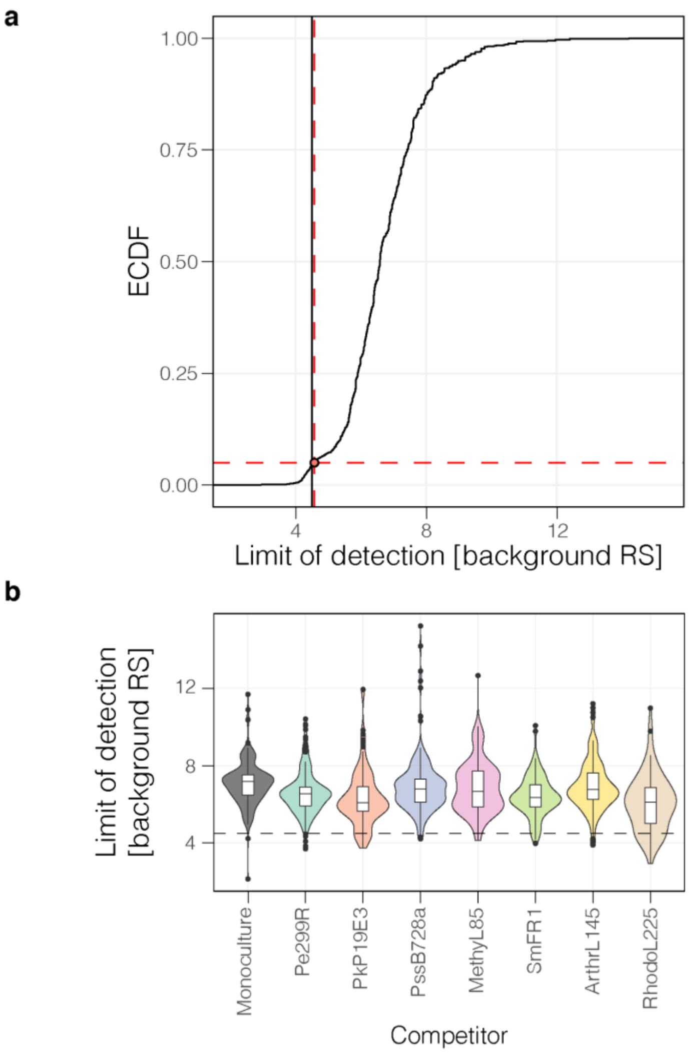


138

139 **Figure S8. Limit of detection of Pe299R$_{CUSPER}$. (a)** Empirical cumulative distribution function (ECDF) of theoretical reproductive success calculated from background fluorescence (background RS). Red dashed lines indicate the 5% threshold of background fluorescent RS ($RS_{LOD}$ = 4.58). Continuous vertical line indicates the limit of detection set at $RS_{>4}$ = 4.50. **(b)** Distribution of background RS of each field of view per treatment group. Dashed line represents the limit of detection.

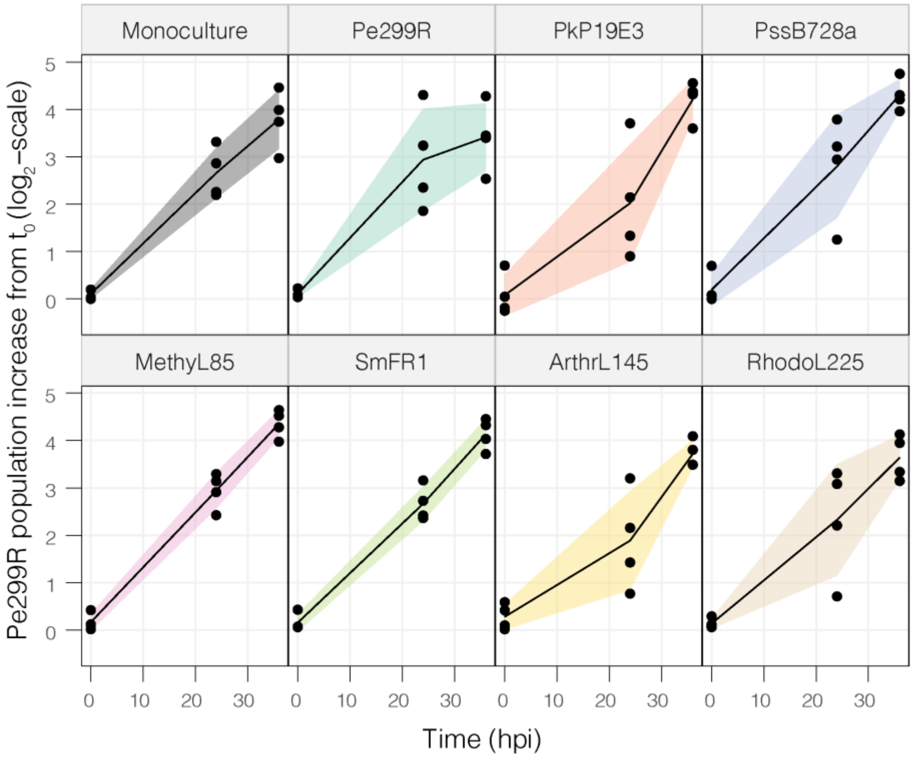


**Figure S9. Increase of *Pe*299R$_{CUSPER}$ population over time in the presence of a competitor in the phyllosphere.** Increase in population size of Pe299R$_{CUSPER}$ from single-cell fluorescent measurements at 0, 24, and 36 h in relation to the founder Pe299R population (population at time zero, $t_0$). Arabidopsis plants were co-inoculated with Pe299R$_{CUSPER}$ and a second epiphyte (top label). Each point represents the population increase in samples taken from independent plants.

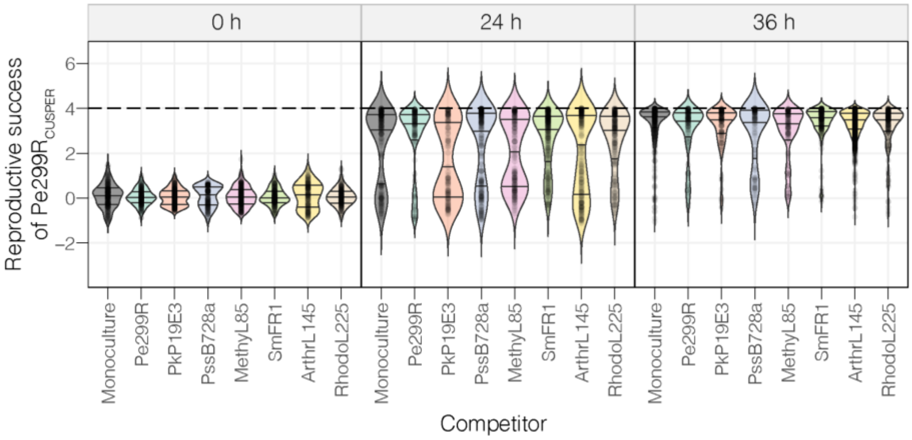


**Figure S10. Single-cell reproductive success of Pe299R$_{CUSPER}$ populations in competition in the phyllosphere.** Distribution of the reproductive success of single Pe299R$_{CUSPER}$ cells in the presence of an epiphyte or as monoculture at 0, 24, and 36 h post-inoculation onto arabidopsis leaves. Each violin plot indicates the median and the interquartile range.

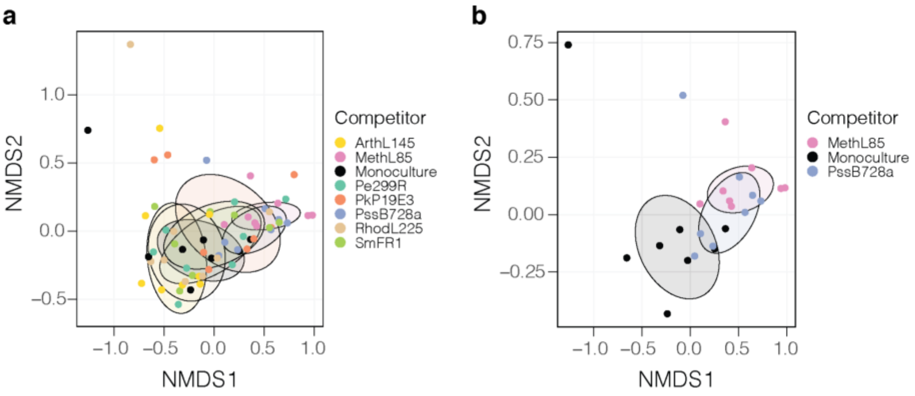


**Figure S11. Non-metric multidimensional analysis of single-cell Pe299R$_{CUSPER}$ population composition in the presence of a second epiphyte. (a)** NMDS was used to discriminate differences in population composition of Pe299R$_{CUSPER}$ at different sampling points and second epiphytes. **(b)** NMDS plot including only significantly different population compositions in relation to the monoculture.

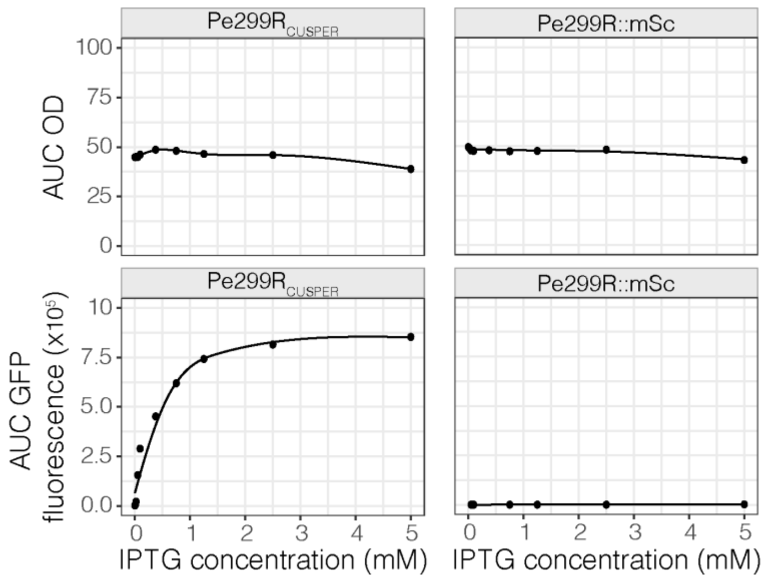


**Figure S12. IPTG concentration curve for Pe299R$_{CUSPER}$ induction.** Optimal concentration of IPTG for the induction of mClover3 in Pe299R$_{CUSPER}$ was evaluated in NB medium supplemented in increasing concentrations of IPTG. Optical density at 600 nm (OD) and green fluorescence were measured over time and the area under the curve (AUC) was used to estimate the biomass and fluorescence yield, respectively, of each growth condition. The parental strain Pe299R::mSc was used as a control.

### SUPPLEMENTAL TABLES

**Table S1.** *In silico* media composition for MRO calculations. Inorganic compounds present in every medium composition are given below.

| Medium | Description | Compound ID (BiGG database) | Compound name |
|---|---|---|---|
| M5C | Minimal medium 5×C | glc__D | D-Glucose |
| | | fru | D-Fructose |
| | | mal__L | L-Malate |
| | | sbt__D | D-Sorbitol |
| | | meoh | Methanol |
| L8C | Leaf-medium 8×C | fum | Fumarate |
| | | mal__L | L-Malate |
| | | glc__D | D-Glucose |
| | | fru | D-Fructose |
| | | glu__L | L-Glutamate |
| | | asp__L | L-Aspartate |
| | | sucr | Sucrose |
| | | meoh | Methanol |
| L10C | Leaf-medium 10×C | fum | Fumarate |
| | | sucr | Sucrose |
| | | asp__L | L-Aspartate |
| | | mal__L | L-Malate |
| | | cit | Citrate |
| | | glu__L | L-Glutamate |
| | | ala__L | L-Alanine |
| | | fru | D-Fructose |
| | | thr__L | L-Threonine |
| | | meoh | Methanol |
| L13C | Leaf-medium 13×C | fum | Fumarate |
| | | lac__L | L-Lactate |
| | | mal__L | L-Malate |
| | | ala__L | L-Alanine |
| | | fru | D-Fructose |
| | | glc__D | D-Glucose |
| | | glu__L | L-Glutamate |
| | | asp__L | L-Aspartate |
| | | chol | Choline |
| | | 4abut | 4-Aminobutanoate (GABA) |
| | | phe__L | L-Phenylalanine |
| | | tyr__L | L-Tyrosine |
| | | meoh | Methanol |
| L18C | Leaf-medium 18×C | fum | Fumarate |
| | | mal__L | L-Malate |
| | | glc__D | D-Glucose |
| | | fru | D-Fructose |
| | | glu__L | L-Glutamate |

| | | | |
|---|---|---|---|
| | | ala__L | L-Alanine |
| | | asp__L | L-Aspartate |
| | | sucr | Sucrose |
| | | gal | D-Galactose |
| | | arab__L | L-Arabinose |
| | | mnl | D-Mannitol |
| | | cit | Citrate |
| | | thr__L | L-Threonine |
| | | gln__L | L-Glutamine |
| | | lys__L | L-Lysine |
| | | asn__L | L-Asparagine |
| | | 4abut | 4-Aminobutanoate (GABA) |
| | | meoh | Methanol |
| L26C | Leaf-medium 26×C | ala__L | L-Alanine |
| | | asp__L | L-Aspartate |
| | | asn__L | L-Asparagine |
| | | chol | Choline |
| | | citr__L | L-Citrulline |
| | | fru | D-Fructose |
| | | fum | Fumarate |
| | | 4abut | 4-Aminobutanoate (GABA) |
| | | glc__D | D-Glucose |
| | | glu__L | L-Glutamate |
| | | gln__L | L-Glutamine |
| | | gly | Glycine |
| | | ile__L | L-Isoleucine |
| | | lac__L | L-Lactate |
| | | leu__L | L-Leucine |
| | | mal__L | L-Malate |
| | | meoh | Methanol |
| | | inost | Myo-Inositol |
| | | orn | Ornithine |
| | | phe__L | L-Phenylalanine |
| | | succ | Succinate |
| | | sucr | Sucrose |
| | | thr__L | L-Threonine |
| | | trp__L | L-Tryptophan |
| | | tyr__L | L-Tyrosine |
| | | val__L | L-Valine |

Inorganic compounds: $Ca^{2+}$, $Cl^{-}$, $Co^{2+}$, $Cu^{2+}$, $Fe^{2+}$, $Fe^{3+}$, $H_2O$, $H^+$, $K^+$, $Mg^{2+}$, $Mn^{2+}$, Molybdate, $Na^+$, Ammonium, $Ni^{2+}$, $O_2$, Phosphate, Sulfate, $Zn^{2+}$.

**Table S2.** Growth parameters of each epiphyte growing in minimal medium supplemented with one (MM + C) or a mix of carbon sources ($MM_{5\times C}$).

| Strain | Growth medium | K (OD) | $\mu$ ($h^{-1}$) |
|---|---|---|---|
| ArthL145 | MM + Fructose | 0.551 | 0.598 |
| | MM + Glucose | 0.515 | 1.382 |
| | MM + Malate | 0.231 | 1.499 |
| | MM + Methanol | 0.000 | 0.000 |
| | MM + Sorbitol | 0.039 | 0.022 |
| | $MM_{5\times C}$ | 0.426 | 0.953 |
| MethL85 | MM + Fructose | 0.029 | 0.026 |
| | MM + Glucose | 0.041 | 0.149 |
| | MM + Malate | 0.401 | 0.157 |
| | MM + Methanol | 0.227 | 0.091 |
| | MM + Sorbitol | 0.067 | 0.056 |
| | $MM_{5\times C}$ | 0.298 | 0.366 |
| Pe299R | MM + Fructose | 0.529 | 1.714 |
| | MM + Glucose | 0.470 | 1.754 |
| | MM + Malate | 0.198 | 1.678 |
| | MM + Methanol | 0.000 | 0.000 |
| | MM + Sorbitol | 0.011 | 0.001 |
| | $MM_{5\times C}$ | 0.419 | 0.937 |
| PkP19E3 | MM + Fructose | 0.564 | 1.486 |
| | MM + Glucose | 0.528 | 0.766 |
| | MM + Malate | 0.245 | 1.141 |
| | MM + Methanol | 0.033 | 0.026 |
| | MM + Sorbitol | 0.021 | 0.002 |
| | $MM_{5\times C}$ | 0.383 | 0.417 |
| PssB728a | MM + Fructose | 0.274 | 1.856 |
| | MM + Glucose | 0.272 | 0.161 |
| | MM + Malate | 0.175 | 0.999 |
| | MM + Methanol | 0.035 | 0.015 |
| | MM + Sorbitol | 0.301 | 0.064 |
| | $MM_{5\times C}$ | 0.301 | 0.959 |
| RhodL225 | MM + Fructose | 0.087 | 1.241 |
| | MM + Glucose | 0.000 | 0.000 |
| | MM + Malate | 0.041 | 2.557 |
| | MM + Methanol | 0.039 | 0.039 |
| | MM + Sorbitol | 0.075 | 0.582 |
| | $MM_{5\times C}$ | 0.317 | 0.709 |
| SmFR1 | MM + Fructose | 0.256 | 1.862 |
| | MM + Glucose | 0.190 | 0.111 |
| | MM + Malate | 0.201 | 0.112 |
| | MM + Methanol | 0.000 | 0.000 |
| | MM + Sorbitol | 0.000 | 0.000 |
| | $MM_{5\times C}$ | 0.328 | 0.751 |

**Table S3.** Growth parameters of Pe299R::mSc from red fluorescence curves in the presence of a competitor or as monoculture.

| Competitor | K (RFU) | μ ($h^{-1}$) | AUC (RFU) | Competition score |
|---|---|---|---|---|
| Monoculture | 1363.69 | 0.26 | 12859.26 | 0.194 |
| ArthL145 | 519.99 | 0.54 | 7586.59 | 2.409 |
| MethL85 | 1041.74 | 0.32 | 11797.01 | 0.924 |
| Pe299R | 663.26 | 0.37 | 8105.26 | 1.947 |
| PkP19E3 | 454.27 | 0.47 | 6172.73 | 2.643 |
| PssB728a | 354.60 | 0.36 | 5129.68 | 3.032 |
| RhodL225 | 1139.11 | 0.34 | 13402.83 | 0.692 |
| SmFR1 | 991.81 | 0.39 | 12181.51 | 1.046 |

**Table S4.** Summary of regression analysis of competitive scores *in vitro*.

| **Model (linear regression)** | ***df*** | **RSE** | **$R^2$** | **$\eta^2$** | ***F*** | ***p*** |
|---|---|---|---|---|---|---|
| Competition score ~ $MRO_{M5C}$ | 22 | 0.70 | 0.461 | 0.484 | 20.7 | 0.0002 |
| **Predictor** | **df** | **SE** | ***b*** | **95% CI** | ***t*** | ***p*** |
| Intercept | 22 | 2.00 | -7.28 | [-11.43, -3.13] | -3.64 | < 0.05 |
| $MRO_{M5C}$ | 22 | 2.87 | 13.04 | [7.092, 18.99] | 4.55 | < 0.05 |

| **Model (linear regression)** | ***df*** | **RSE** | **$R^2$** | **$\eta^2$** | ***F*** | ***p*** |
|---|---|---|---|---|---|---|
| Competition score ~ MRO (without PssB728a) | 18 | 0.36 | 0.81 | 0.823 | 83.45 | < 0.05 |
| **Predictor** | **df** | **SE** | ***b*** | **95% CI** | ***t*** | ***p*** |
| Intercept | 18 | 1.04 | -7.97 | [-10.16, -5.77] | -7.63 | < 0.05 |
| $MRO_{M5C}$ | 18 | 1.49 | 13.65 | [10.51, 16.78] | 9.14 | < 0.05 |

| **Model (GLM, Gamma distribution, log link)** | ***df*** | **SE** | **Null Deviance** | **Residual Deviance** | **Pseudo-$R^2$** | ***F*** | ***p*** |
|---|---|---|---|---|---|---|---|
| Competition score ~ MRO × PD | 20 | 0.0424 | 7.60 | 0.801 | 0.89 | 9.61 | 0.0056 |
| **Predictor** | **df** | **SE** | ***b*** | **95% CI** | ***t*** | ***p*** | |
| Intercept | 20 | 4.10 | 8.88 | [0.98, 16.74] | 2.17 | 0.043 | |
| $MRO_{M5C}$ | 20 | 5.73 | -9.90 | [-20.89, 1.18] | -1.73 | 0.099 | |
| PD | 20 | 5.64 | -19.19 | [-29.93, -8.40] | -3.41 | 0.0028 | |
| $MRO_{M5C}$ × PD | 20 | 7.87 | 24.06 | [8.92, 39.076] | 3.06 | 0.0062 | |

**Table S5.** Summary of regression analysis of competitive scores *in planta*.

| Model (y ~ x) | *df* | RSE | $R^2$ | *F* | *p* |
|---|---|---|---|---|---|
| Competition score ~ PD | 4 | 1.09 | -0.19 | 0.20 | 0.6763 |
| Competition score ~ PD × $MRO_{Complete}$ | 2 | 1.05 | -0.11 | 0.84 | 0.5835 |
| Competition score ~ PD × $MRO_{M5C}$ | 2 | 0.91 | 0.17 | 1.33 | 0.4556 |
| Competition score ~ PD × $MRO_{L8C}$ | 2 | 1.15 | -0.32 | 0.60 | 0.6747 |
| **Competition score ~ PD × $MRO_{L10C}$** | **2** | **0.28** | **0.92** | **20.15** | **0.0477** |
| Competition score ~ PD × $MRO_{L13C}$ | 2 | 1.05 | -0.10 | 0.85 | 0.5809 |
| Competition score ~ PD × $MRO_{L18C}$ | 2 | 1.20 | -0.43 | 0.50 | 0.7202 |
| Competition score ~ PD × $MRO_{L26C}$ | 2 | 1.13 | -0.27 | 0.65 | 0.6543 |

| Predictor (Competition score ~ PD × $MRO_{L10C}$) | *df* | SE | *b* | 95% CI | $\eta^2$ | *t* | *p* |
|---|---|---|---|---|---|---|---|
| Intercept | 2 | 19.14 | -114.05 | [-196.42, -31.68] | | -5.96 | 0.027 |
| $MRO_{L10C}$ | 2 | 29.29 | 176.62 | [50.61, 302.63] | 0.390 | 6.03 | 0.026 |
| PD | 2 | 25.44 | 142.94 | [33.49, 252.39] | 0.170 | 5.62 | 0.030 |
| $MRO_{L10C}$ × PD | 2 | 38.90 | -222.77 | [-390.15, -55.39] | 0.530 | -5.73 | 0.029 |

**Table S6.** Summary of PERMANOVA on Pe299$R_{CUSPER}$ populations.

| Model | Term | *df* | SS | $R^2$ | *F* | *p* |
|---|---|---|---|---|---|---|
| Full Model | Time | 1 | 0.585 | 0.100 | 8.695 | 0.001 |
| | Competitor | 7 | 1.583 | 0.270 | 3.363 | 0.002 |
| | Residual | 55 | 3.699 | 0.630 | n.a. | n.a. |
| | Total | 63 | 5.868 | 1.000 | n.a. | n.a. |
| Monoculture vs PssB728a | Competitor | 1 | 0.324 | 0.243 | 4.498 | 0.011 |
| | Residual | 14 | 1.008 | 0.757 | n.a. | n.a. |
| | Total | 15 | 1.331 | 1.000 | n.a. | n.a. |
| Monoculture vs MethL85 | Competitor | 1 | 0.532 | 0.358 | 7.802 | 0.003 |
| | Residual | 14 | 0.954 | 0.642 | n.a. | n.a. |
| | Total | 15 | 1.486 | 1.000 | n.a. | n.a. |
| 24 h vs 36 h | Time | 1 | 0.585 | 0.100 | 6.864 | 0.003 |
| | Residual | 62 | 5.283 | 0.900 | n.a. | n.a. |
| | Total | 63 | 5.868 | 1.000 | n.a. | n.a. |

**Table S7.** Primers used to construct pProbe_CUSPER.

| **Name** | **Target** | **Sequence (5'-3')[a]** | ***tm* (°C)[b]** |
|---|---|---|---|
| Plac_fw | $P_{A1/04/03}$ promoter | tcctcgcccttgctcatAAATTGTTATCCGCTCACAATTG | 55 |
| Plac_rv | $P_{A1/04/03}$ promoter | tgccactcatcgcagtactGAAAATTTATCAAAAAGAGTG | 47 |
| fp_fw | *mClover3* gene | tgagcggataacaatttATGAGCAAGGGCGAGGAGCTG | 64 |
| fp_rv | *mClover3* gene | actggaaagcgggcagtgaATTCTCACCAATAAAAAACG | 50 |
| lacIq_fw | *lacIq* repressor gene | aagtaccgccacctaaGACACCATCGAATGGTGCAAAACC | 61 |
| lacIq_rv | *lacIq* repressor gene | gttttttattggtgagaatTCACTGCCCGCTTTCCAGTCG | 62 |
| gmR_fw | *gmR* selection gene | aggaattggggatcggaagcttTTGACATAAGCCTGTTCGG | 53 |
| gmR_rv | *gmR* selection gene | ccattcgatggtgtcTTAGGTGGCGGTACTTGGGTCG | 61 |

[a] Capital letters indicate complementary sequences to the original PCR target; lower cases indicate complementary sequences to the destination sequence (overhang).

[b] The *tm* refers to the melting temperature of the original target sequence.
